# Ubiquitin-dependent and -independent roles of E3 ligase RIPLET in innate immunity

**DOI:** 10.1101/367656

**Authors:** Cristhian Cadena, Sadeem Ahmad, Audrey Xavier, Joschka Willemsen, Sehoon Park, Ji Woo Park, Seong-Wook Oh, Takashi Fujita, Fajian Hou, Marco Binder, Sun Hur

## Abstract

The conventional view posits that E3 ligases function primarily through conjugating ubiquitin (Ub) to their substrate molecules. We report here that RIPLET, an essential E3 ligase in antiviral immunity, promotes the antiviral signaling activity of the viral RNA receptor RIG-I through both Ub-dependent and -independent manners. RIPLET uses its dimeric structure and a bivalent binding mode to preferentially recognize and ubiquitinate RIG-I pre-oligomerized on dsRNA. In addition, RIPLET can cross-bridge RIG-I filaments on longer dsRNAs, inducing aggregate-like RIG-I assemblies. The consequent receptor clustering synergizes with the Ub-dependent mechanism to amplify RIG-I-mediated antiviral signaling in an RNA-length dependent manner. These observations show the unexpected role of an E3 ligase as a co-receptor that directly participates in receptor oligomerization and ligand discrimination. It also highlights a previously unrecognized mechanism by which the innate immune system measures foreign nucleic acid length, a common criterion for self vs. non-self nucleic acid discrimination.

## INTRODUCTION

Effective immune response to viral infection depends on the efficient recognition of viral RNAs by an innate immune receptor, RIG-I. RIG-I detects a broad range of viral RNAs, by recognizing double-stranded RNA (dsRNA) harboring a 5’-triphosphate (5’ppp) or diphosphate group (Goubau et al., 2014; Schlee et al., 2009). These features are often present in viral RNAs, either in the form of defective interfering particles or in viral genomes, but are typically absent in cellular RNAs. Viral RNA recognition then leads to the activation of the downstream adaptor molecule, MAVS, which triggers antiviral signaling pathways to produce type I and III interferons (IFNs) and other inflammatory cytokines (Yoneyama and Fujita, 2010).

Previous structural and biochemical studies have revealed a detailed picture of how RIG-I recognizes dsRNA with 5’ppp and how viral RNA binding leads to signal activation (Lassig and Hopfner, 2017; Rawling and Pyle, 2014; Sohn and Hur, 2016). In the absence of viral RNA, RIG-I is in an auto-repressed state wherein the signaling domain, the N-terminal tandem CARD (2CARD), is inhibited from activating MAVS (Kowalinski et al., 2011). dsRNA binding occurs via the RIG-I helicase domain and the C-terminal domain (CTD), which together bind the dsRNA end and recognize the 5’ppp (Jiang et al., 2011; Luo et al., 2012). dsRNA binding and/or the subsequent ATP binding was proposed to release the auto-repression of 2CARD (Kowalinski et al., 2011; Rawling et al., 2015). The released 2CARD then forms a tetramer and recruits MAVS through a homotypic CARD-CARD interaction (Jiang et al., 2012; Peisley et al., 2014). This interaction nucleates MAVS CARD filament formation, which in turn serves as a signaling platform to recruit and activate downstream signaling molecules, such as TBK1 and IRF3 (Hou et al., 2011; Wu et al., 2014).

The notion that RIG-I signaling requires not only a release of 2CARD but also its tetramerization is supported by the presence of multiple mechanisms that tightly regulate the 2CARD tetramerization process. First, while RIG-I binds to dsRNA ends as a monomer, it can also form filamentous oligomers on longer dsRNA through ATP-driven translocation of individual monomers from the dsRNA end to the interior (Binder et al., 2011; Devarkar et al., 2018; Myong et al., 2009; Patel et al., 2013; Peisley et al., 2013). It was further shown that filament formation of RIG-I promotes 2CARD tetramerization, presumably by increasing the local concentration of 2CARD (Peisley et al., 2013). Second, in addition to RIG-I filament formation, K63-linked polyubiquitin (K63-Ub_n_) was also shown to promote RIG-I signaling (Jiang et al., 2012). Structural studies further showed that K63-Ub_n_ chains bind the periphery of the core 2CARD tetramer, bridging the adjacent subunits and stabilizing its assembly (Peisley et al., 2014). The 2CARD tetramer then acts as a helical template to nucleate the MAVS filament formation for downstream signal activation (Wu and Hur, 2015).

Despite the detailed understanding of the action of K63-Ub_n_ on RIG-I, much remains debated as to how and when Ub is placed on RIG-I, which E3 ligase is involved and how it interplays with RIG-I filament formation. Previous studies reported that TRIM25 and RIPLET (*i.e.* RNF135) are two essential E3 ligases important for RIG-I signaling (Gack et al., 2007; Gao et al., 2009; Oshiumi et al., 2009; Oshiumi et al., 2010). A more recent study proposed that RIPLET and TRIM25 sequentially act on RIG-I upon viral RNA engagement (Oshiumi et al., 2013). That is, RIPLET first acts on the C-terminal portion of RIG-I, releasing 2CARD, which is then modified by TRIM25. However, given the previous finding that RNA binding is sufficient to release 2CARD (Kowalinski et al., 2011), it was unclear whether RIPLET indeed acts to release 2CARD and if so, how. At the same time, accumulating evidence suggested that TRIM25 has multiple, RIG-I-independent antiviral functions (Choudhury et al., 2014; Li et al., 2017; Manokaran et al., 2015; Meyerson et al., 2017; Zheng et al., 2017), raising the question whether the observed effect of TRIM25 on RIG-I represents a direct or an indirect effect.

We here report a combination of cellular and biochemical data showing that RIG-I activation is dependent on RIPLET, not TRIM25, and that RIPLET suffices to ubiquitinate and activate RIG-I. In addition, RIPLET recognizes the filamentous form of RIG-I using two distinct binding modes, the interplays of which offer previously unrecognized mechanisms for ligand discrimination and receptor clustering.

## RESULTS

### RIPLET, not TRIM25, is required for RIG-I signaling

To examine the importance of RIPLET and TRIM25 in RIG-I functions, we first measured the signaling activity of RIG-I in wild-type (WT) 293T cells and those lacking RIG-I, RIPLET or TRIM25 (Shi et al., 2017). Cells were stimulated with either *in vitro* transcribed 42 bp dsRNA with 5’ppp (Figure 1A) or Sendai virus (SeV) (Figure 1B), stimuli known to trigger RIG-I signaling. Comparison of the level of mRNAs for IFNβ or IFN-stimulated genes (ISGs) showed that RIG-I and RIPLET are required for antiviral signaling in response to 5’ppp-dsRNA and SeV. TRIM25, however, was not required as TRIM25^−/−^ cells displayed slightly enhanced antiviral signaling. Similar activities were observed with the IFNβ promoter driven luciferase activity assay (Figure 1C). RIPLET^−/−^ 293T cells were normal in MDA5- and MAVS-mediated signaling (Figure 1D), suggesting that the defect in antiviral signaling is specific to RIG-I. Impaired RIG-I signaling in RIPLET^−/−^ 293T was restored by complementation with RIPLET (Figure 1E), further supporting the notion that RIPLET, not TRIM25, is required for RIG-I signaling.

**Fig. 1.**
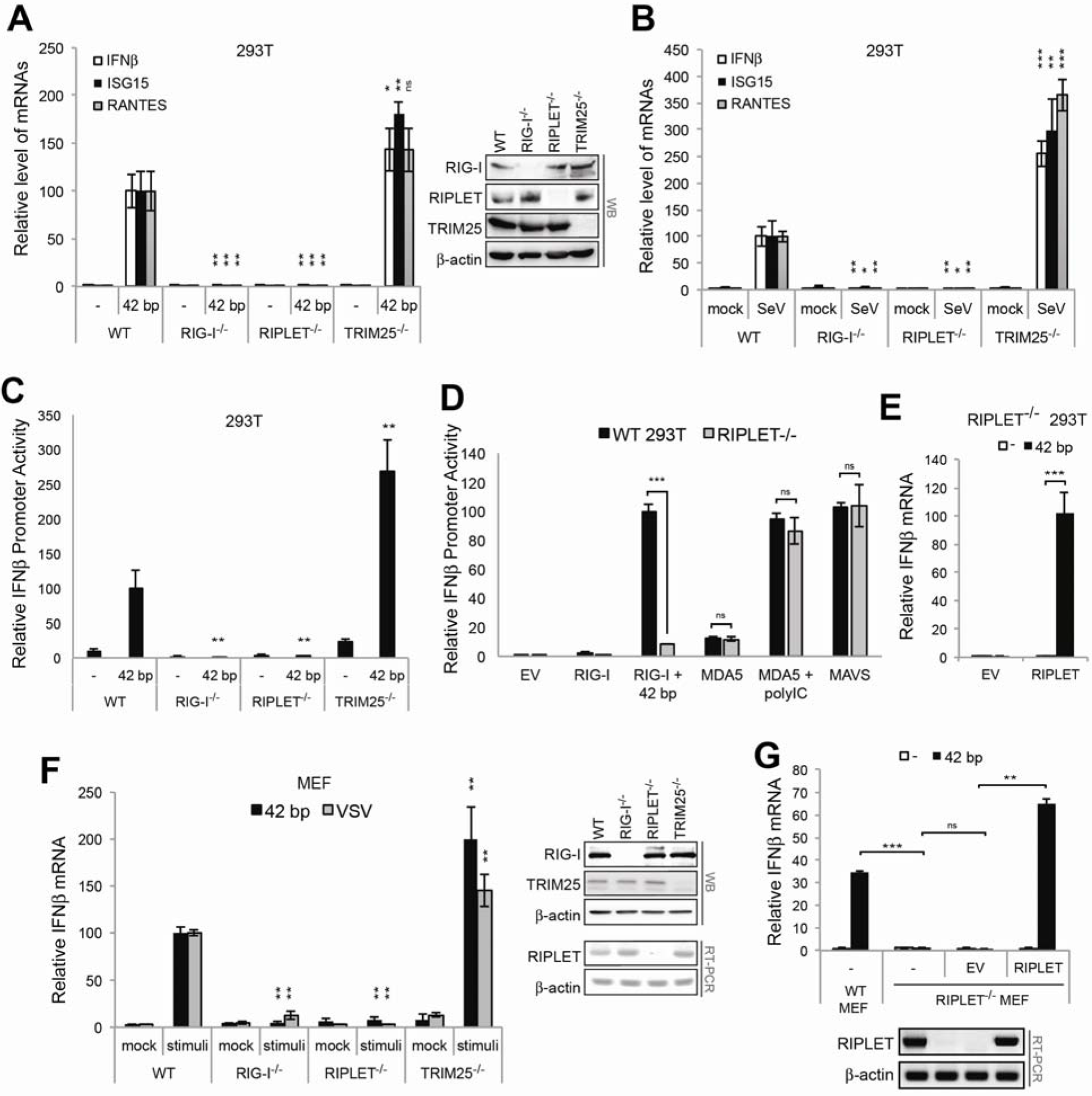
RIPLET, not TRIM25, is required for RIG-I signaling. A-B. Relative level of antiviral signaling in 293T cells (WT, RIG-I^−/−^, RIPLET^−/−^ or TRIM25^−/−^), as measured by the level of IFNβ ISG15 and RANTES mRNAs. Cells were stimulated with 42 bp dsRNA with 5’ppp (0.2 μg) (A) or SeV (100 HA units/ml). Unless mentioned otherwise, all RNAs used in this manuscript harbor 5’ppp. The level of mRNAs were normalized against the WT level with 42 bp dsRNA. Right of (A): western blot (WB) showing deletion of respective genes. C. Relative level of antiviral signaling in WT, RIG-I^−/−^, RIPLET^−/−^ or TRIM25^−/−^ 293T cells, as measured by the IFNβ promoter reporter assay. Cells were stimulated with 42 bp dsRNA as in (A). D. Relative level of RIG-I, MDA5 and MAVS signaling in RIPLET^−/−^ 293T cells. Cells were transfected with plasmid expressing RIG-I (10 ng), MDA5 (5 ng) or MAVS (25 ng), and were stimulated with 42 bp dsRNA (0.2 μg) or high molecular weight polyinosinic-polycytidylic acid (polyIC, 0.5 μg). Note that ectopic expression of MDA5 and MAVS is necessary to examine their signaling activities in 293T cells. RIG-I was ectopically expressed for comparison. The rest of Figure 1 measures the endogenous RIG-I activity. E. Relative level of antiviral signaling in RIPLET^−/−^ 293T cells with and without RIPLET complementation. 293T cells were transfected with plasmid expressing RIPLET (or empty vector, EV, 10 ng) and stimulated with 42 bp dsRNA. F. Relative level of antiviral signaling in MEF cells (WT, RIG-I^−/−^, RIPLET^−/−^ or TRIM25^−/−^), as measured by the level of IFNβ mRNA. Cells were stimulated with 42 bp dsRNA with 5’ppp (0.2 μg) or VSV (MOI=1). The level of mRNAs were normalized against the WT level with 42 bp dsRNA. Right: WB or RT-PCR analysis showing deletion of respective genes. For RIPLET, mRNA level was measured due to the lack of an appropriate antibody against mouse RIPLET. Note that the αRIPLET WB in (A) is for human RIPLET. G. Relative level of antiviral signaling in RIPLET^−/−^ MEF cells with and without stable expression of RIPLET using lentivirus transduction. All data are presented as mean ± s.d. (*n* = 3-4) and are representative of at least three independent experiments. * *p* < 0.05, ** *p* < 0.01, \*\*\**p* < 0.001, ns, not significant (unpaired *t* test). Unless indicated otherwise, *p* values were calculated in comparison to WT cell values. All cell lines used in this figure (except for (G), which is from (Oshiumi et al., 2010), see Figure S1A) were from (Shi et al., 2017).

The importance of RIPLET and the non-essential role of TRIM25 in RIG-I signaling was also observed with mouse embryonic fibroblasts (MEFs). Stimulation of MEFs with 5’ppp-dsRNA or vesicular stomatitis virus (VSV) triggered antiviral signaling as measured by the induction of IFNβ mRNA (Figure 1F). This antiviral signaling activity depended on RIG-I and RIPLET, but not TRIM25, in agreement with the previous report (Shi et al., 2017). Complementation with RIPLET restored the antiviral signaling activity in RIPLET^−/−^ MEFs (Figure 1G).

To further validate our observation, we examined additional knock-out cell lines from independent sources (Figures S1A-S1E). These included RIPLET^−/−^ MEF (from (Oshiumi et al., 2010)), TRIM25^−/−^ 293T and A549 (from the Binder lab, see STAR methods) and TRIM25^−/−^ MEF (from (Gack et al., 2007)). 5’ppp-dsRNA, SeV, VSV, Rift Valley fever virus (RVFV) and influenza A virus (FluA) were used to stimulate RIG-I signaling. To ensure that the lack of dependence of RIG-I signaling on TRIM25 is not due to particular experimental conditions, viral assays with A549 cells (Figures S1D & S1E) were independently performed in a different laboratory using different protocols (see STAR methods). The dependence of RIG-I on RIPLET was further confirmed in independently generated RIPLET^−/−^ MEF cell line (Figure S1A). A moderate reduction (~40 %) in RIG-I signaling was observed only with one of the TRIM25^−/−^ MEF cell lines (Figure S1C), while all other TRIM25^−/−^ cells showed little defect in RIG-I signaling (Figures 1, S1B, S1D & S1E). TRIM25^−/−^ cells also displayed no defect in MDA5 signaling (Figures S1F & S1G)

Previous studies suggested that TRIM25 is required for the signaling activity of isolated 2CARD (Gack et al., 2007). In our analysis, TRIM25 knock-out led to ~50 % reduction in the signaling activity of GST-2CARD (Figure S1H), and ectopic expression of TRIM25 partially restored it (Figure S1I). This result suggests that while TRIM25 is not necessary for full-length RIG-I signaling, it has a moderate stimulatory effect on the signaling activity of the isolated 2CARD. Note that the signaling activity of GST-2CARD was unaffected by RIPLET deletion (Figure S1H). The observed gain of TRIM25-dependence and the loss of RIPLET-dependence for GST-2CARD suggest that the signaling mechanism of GST-2CARD differs from that of full-length RIG-I.

### RIPLET, not TRIM25, ubiquitinates RIG-I in a dsRNA-dependent manner

To test whether RIPLET or TRIM25 can directly conjugate K63-Ub_n_ to RIG-I, we reconstituted the RIG-I ubiquitination reaction *in vitro*. A previous study showed that RIG-I signaling requires Ubc13 (*i.e.* Ube2N) (Shi et al., 2017), an Ub E2 conjugating enzyme that can function either by itself or in complex with catalytically inactive E2 variants, such as Uev1A (*i.e.* Ube2V1) (Wu and Karin, 2015). Incubation of RIG-I with purified E1, E2(s), and RIPLET showed that the Ubc13:Uev1A complex, but not Ubc13 alone, supported robust ubiquitination of RIG-I (Figure 2A). Consistent with the lack of effect of RIPLET on MDA5 signaling, MDA5 was not ubiquitinated by RIPLET under the same condition (Figure S2A). The K63R mutation in Ub prevented polyUb chain conjugation by RIPLET (Figure 2B), suggesting that RIPLET primarily conjugates K63-Ub_n_ to RIG-I. Although ubiquitinated RIG-I migrated into the SDS gel as a discrete band, it represents polyubiquitinated RIG-I (likely compressed in the resolving and stacking gel interface). This is evidenced by the appearance of smaller species when RIG-I was ubiquitinated with an increasing amount of the K63 chain terminator, K63R Ub (Figure 2C). Intriguingly, ubiquitination of RIG-I required presence of dsRNA (Figure 2A), recapitulating the cellular requirement of dsRNA for RIG-I signaling. Since RIPLET does not appear to bind dsRNA (Figure S2B), dsRNA-dependent ubiquitination suggests the specificity of RIPLET for the RIG-I:dsRNA complex over RNA-free RIG-I (to be confirmed in Figure 3).

**Fig. 2.**
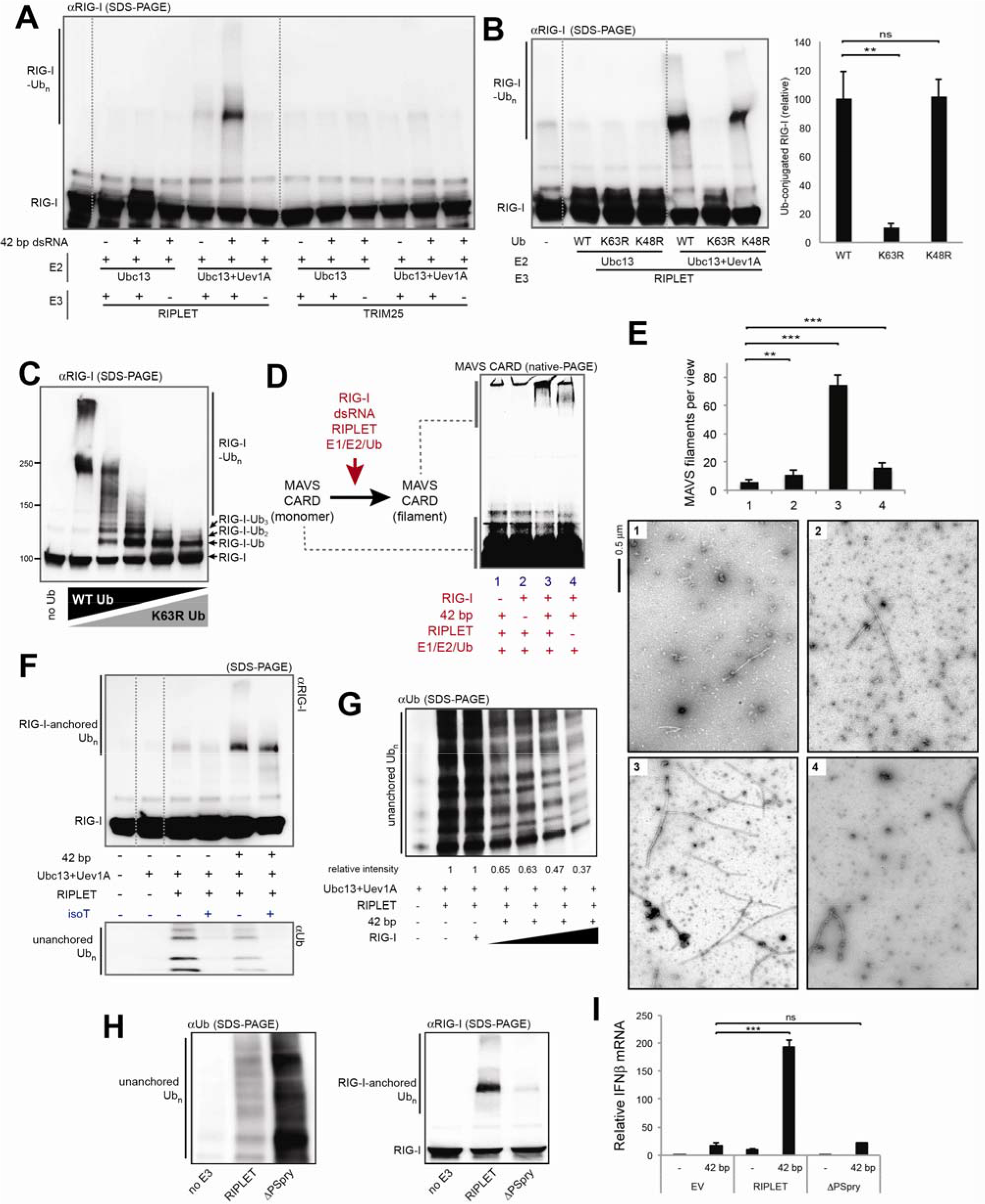
RIPLET, not TRIM25, ubiquitinates RIG-I in a dsRNA-dependent manner. A. *In vitro* ubiquitination analysis of RIG-I. Purified E1 (1 μM), E2 (Ubc13 alone or in complex with Uev1A, 5 and 2.5 μM, respectably), E3 (RIPLET or TRIM25, 0.25 μM) and Ub (20 μM) were incubated with RIG-I (0.5 μM), either alone or in complex with 42 bp dsRNA (1 ng/μl). Ubiquitination of RIG-I was analyzed by anti-RIG-I blot on SDS gels. B. Ub linkage type analysis. RIG-I was ubiquitinated by RIPLET using Ub (WT, K63R or K48R) in the presence of 42 bp dsRNA. Right: quantitation of ubiquitinated RIG-I (mean ± s.d., *n* = 3). Ubiquitination reactions were performed as in (A). * *p* < 0.05, ** *p* < 0.01, \*\*\**p* < 0.001, ns, not significant (unpaired *t* test, compared with WT Ub). C. *In vitro* ubiquitination analysis of RIG-I in the presence of an increasing amount of K63R Ub (0, 25, 50, 75, 100%) mixed in with WT Ub. Total Ub concentration (20 μM) was kept constant. All reactions were performed in the presence of 42 bp dsRNA. D-E. RIG-I- and RIPLET-induced MAVS polymerization assay. RIG-I was ubiquitinated by RIPLET as in (A), and was mixed with purified, monomeric MAVS CARD fused to a SNAP tag (10 μM). Monomer-to-filament transition of MAVS CARD was analyzed by native gel assay (D) or negative stain EM (E). For native gel analysis, fluorescence of Alexa647 conjugated to the SNAP tag was used for gel imaging. For the EM analysis, six random images (representative images shown below) were collected at 4,800x magnification, and average number (± s.d) of filaments per image was plotted. * *p* < 0.05, ** *p* < 0.01, \*\*\**p* < 0.001, ns, not significant (unpaired *t* test, compared with lane 1). F. Synthesis of anchored and unanchored Ub chains by RIPLET. The ubiquitination reaction was performed as in (A). Anchored and unanchored Ub chains were analyzed by anti-RIG-I and anti-Ub blotting, respectively. Only the low molecular weight species (<~50 kDa) were analyzed for anti-Ub blot to avoid detection of RIG-I-conjugated Ub, which would be > ~100 kDa. IsoT (10 ng/μl) was used to further confirm that the lower molecular weight species are unanchored Ub chains. Note that RIG-I-anchored Ub chains were also slightly degraded by isoT. G. Unanchored Ub chain synthesis by RIPLET, with and without RIG-I (0.5 μM for lane 3 and 0.5-1.5 μM for lanes 4-7) and 42 bp dsRNA (3 ng/μl). The reaction was performed as in (A), and analyzed by anti-Ub blot as in (F). Averaged relative intensity of unanchored Ub chain from triplicate experiments were indicated below the image (s.d.<0.05). H. RIG-I-anchored vs. unanchored Ub chain synthesis by RIPLET and ΔPSpry. Unanchored Ub synthesis was measured in the absence of RIG-I or dsRNA. RIG-I-anchored Ub synthesis was measured in the presence of RIG-I and 42 bp dsRNA. All reactions were performed using conditions equivalent to (A). I. Relative level of antiviral signaling (mean ± s.d., *n* = 3) in RIPLET^−/−^ 293T cells when complemented with empty vector (EV), RIPLET or ΔPSpry (10 ng plasmid). * *p* < 0.05, ** *p* < 0.01, \*\*\**p* < 0.001, ns, not significant (unpaired *t* test). All data are representative of at least three independent experiments.

**Fig. 3.**
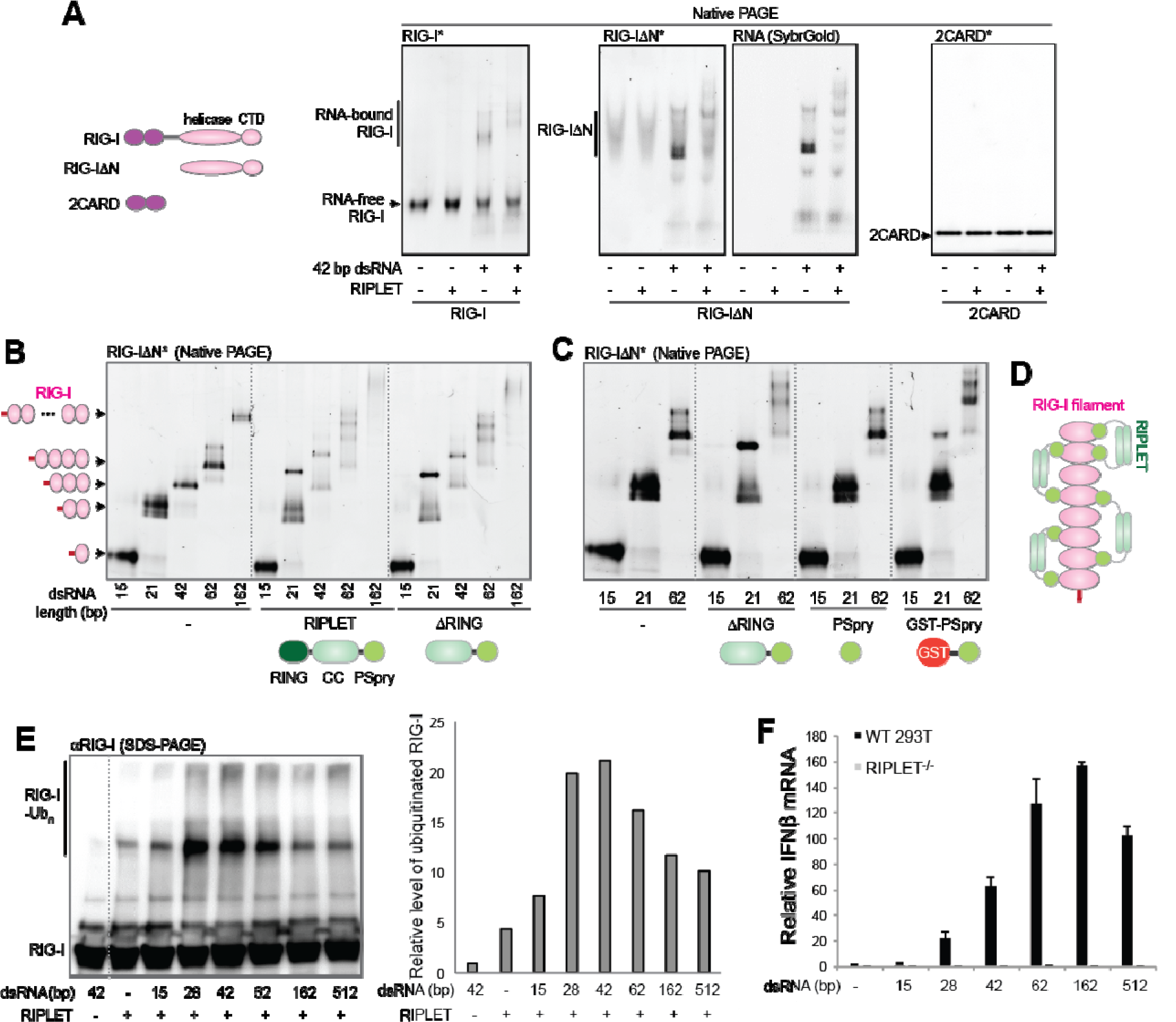
RIPLET recognizes pre-oligomerized RIG-I on dsRNA and activates RIG-I signaling in a dsRNA length-dependent manner. a. Native gel shift assay to monitor the interaction between RIG-I and RIPLET. Fluorescently labeled RIG-I (full-length RIG-I, RIG-IΔN or isolated 2CARD, 0.5 μM) was incubated with RIPLET (0.5 μM) in the presence or absence of 42 bp dsRNA (1 ng/μl). The complex was analyzed by native gel using RIG-I fluorescence (*). For RIG-IΔN, SybrGold stained image was also shown to clearly visualize only the RNA-bound species. B-C. Native gel shift assay of RIG-IΔN (0.5 μM) with and without RIPLET (B), ΔRING (B, C), PSpry (C) or GST-PSpry (C) (0.5 μM for all RIPLET variants). 15-162 bp dsRNAs (1 ng/μl) were used to form the RIG-I:dsRNA complexes of various sizes. Gel images were acquired using RIG-I fluorescence (*). D. A model of how RIPLET binds RIG-I filaments. Based on the dsRNA length dependence and bivalency requirement, we propose that RIPLET binds two nearby RIG-I monomers within a filament (either immediately adjacent or separated). The lengths of RIPLET CC (~60 amino acids, expected to be ~80 Å) and the linker between CC and PSpry (~50 amino acids, expected to be >100 Å for fully disordered linker) are compatible with the model. E. *In vitro* ubiquitination analysis of RIG-I in the presence of 15-512 bp dsRNAs (1 ng/μl for all RNAs), as performed in Figure 2A. Right: densitometry analysis of ubiquitinated RIG-I, representative of three independent analyses. Anti-RIG-I immunoblot was used for quantitation. A similar dsRNA length-dependence was observed by fluorescence measurement using FITC-labeled RIG-I. F. Relative level of antiviral signaling in response to 15-512 bp dsRNA (0.2 μg for all RNAs) in 293T cells (WT or RIPLET^−/−^), as measured by the IFNβ mRNA level (mean ± s.d., *n* = 3). See Figure S3H for the comparison between WT and RIG-I^−/−^ 293T cells. All data are representative of at least three independent experiments.

Consistent with the non-essential role of TRIM25 for RIG-I signaling (Figures 1 and S1), no RIG-I ubiquitination was observed with TRIM25 under the equivalent condition (Figure 2A). While the lack of ubiquitination could be due to the use of inappropriate E2 enzymes, the Ubc13:Uev1A complex was found required for RIG-I signaling (Shi et al., 2017) and is so far the only cytosolic E2 known to synthesize K63-linked Ub chains, the linkage type required for 2CARD tetramerization (Peisley et al., 2014; Zeng et al., 2010). Under the same condition, TRIM25 robustly synthesized unanchored Ub chains (Figure S2C) and auto-ubiquitinated itself (Figure S2D), suggesting that purified TRIM25 is active. Furthermore, both unanchored Ub synthesis and auto-ubiquitination by TRIM25 showed strong dsRNA dependence (Figures S2C & S2D), which is in line with the reported dsRNA binding activity of TRIM25 (Choudhury et al., 2017; Manokaran et al., 2015). TRIM25 did not ubiquitinate RIG-I with or without dsRNA (Figure 2A). While multiple previous studies, including our own (Peisley et al., 2014), showed *in vitro* ubiquitination of isolated 2CARD by TRIM25, this was done using excessively high protein concentration (e.g. ~10 fold higher TRIM25 and substrate (Peisley et al., 2014)). No study, to our knowledge, has shown a dsRNA-dependent ubiquitination of RIG-I by TRIM25, an activity expected for the E3 ligase that stimulates RIG-I only upon viral infection.

To examine whether RIPLET-mediated Ub-conjugation can activate RIG-I, we employed the *in vitro* MAVS stimulation assay in which the activation of MAVS was examined by its transition from a monomeric to a filamentous state (Peisley et al., 2013; Wu et al., 2013). In this assay, RIG-I was first ubiquitinated by RIPLET, and then was incubated with isolated, monomeric MAVS CARD. The monomer-to-filament transition of MAVS CARD was then measured by native gel electrophoresis (see STAR Methods) or electron microscopy (EM). Note that MAVS filaments are significantly longer (typically >0.1 μm) than RIG-I filaments on 42 bp dsRNA (limited to ~10 nm). This difference allows unambiguous identification of MAVS filaments and quantitative analysis of MAVS filament frequencies (Figure 2E). Both EM and native gel assay showed that MAVS filamentation is slightly promoted by RIG-I in the presence of either dsRNA or RIPLET, but the two together significantly enhance the MAVS-stimulatory activity of RIG-I (Figures 2D & 2E). This is consistent with the cellular signaling activity of RIG-I, which requires both dsRNA and RIPLET (Figure 1).

Previous studies suggested that RIG-I activation is largely dependent on unanchored Ub chains (Shi et al., 2017; Zeng et al., 2010). We first examined whether RIPLET synthesizes unanchored Ub_n_ chains, in addition to the anchored chains, and how unanchored Ub_n_ synthesis is affected by dsRNA. In agreement with the previous report (Shi et al., 2017), we found that RIPLET indeed promotes unanchored Ub_n_ synthesis (Figure 2F). However, this activity was more robust in the absence of RIG-I and was significantly suppressed with an increasing concentration of the RIG-I:dsRNA complex (Figure 2G). This suggests that unanchored Ub_n_ synthesis occurs constitutively, while the RIG-I:dsRNA complex binding triggers transition from unanchored Ub_n_ synthesis to RIG-I-anchored Ub_n_ synthesis. Consistent with this notion, the C-terminal Pry/Spry domain (PSpry, also known as B30.2) of RIPLET, which is required for interaction with RIG-I (to be shown in Figure 3), was not necessary for unanchored Ub_n_ synthesis (Figure 2H). In fact, deletion of PSpry improved the unanchored Ub_n_ synthesis activity. By contrast, the PSpry domain was required for both efficient ubiquitination of RIG-I (Figure 2H) and RIG-I signaling (Figure 2I). In addition, anchored Ub_n_ alone appears to be sufficient for RIG-I signaling; RIG-I maintained a significant level of the MAVS polymerization activity even after treatment with isoT (Figure S2E), an ubiquitin-processing protease with a known preference for unanchored Ub_n_. Although isoT moderately reduced the MAVS polymerization activity, this could be due to partial degradation of anchored Ub_n_ by isoT (as observed in Figure 2F). Consistent with the notion that RIPLET-mediated Ub conjugation can activate RIG-I, mass spectrometric analysis showed that RIPLET ubiquitinates RIG-I at multiple sites, including K190 and K193 on 2CARD, that are located in close proximity to the Ub binding site (Figure S2F, see also Figure S2G). Altogether, our data show that RIPLET is an essential E3 ligase that ubiquitinates RIG-I in a dsRNA-dependent manner. Moreover, RIPLET-mediated ubiquitination is sufficient to activate RIG-I. While RIPLET can stimulate unanchored Ub_n_ synthesis in the absence of RIG-I, this activity appears neither necessary for nor correlated with RIG-I signaling.

### RIPLET recognizes pre-assembled RIG-I filaments on dsRNA

We next explored how RIPLET interacts with RIG-I and how this interaction depends on dsRNA. We purified recombinant RIPLET and RIG-I (Figure S3A) and used native gel shift assays to examine the mobility of fluorescently labeled RIG-I with and without RIPLET. Consistent with the dsRNA-dependent ubiquitination of RIG-I (Figure 2A), RIPLET bound RIG-I only in the presence of dsRNAs (Figure 3A). RIPLET did not bind MDA5 filaments (Figure S3B), consistent with its lack of a role in MDA5 signaling (Figure 1D) and ubiquitination (Figure S2A). The N-terminal 2CARD deletion mutant of RIG-I (RIG-IΔN) also bound RIPLET in a manner dependent on dsRNA. Isolated 2CARD did not bind RIPLET (Figure 3A), in agreement with the lack of effect of RIPLET on GST-2CARD signaling (Figure S1H). Further truncation of RIG-I to the helicase domain or CTD impaired RIPLET binding (Figure S3C), indicating that RIG-IΔN is minimally required. We then compared RIPLET binding to RIG-I in complex with 15, 21, 42, 62, and 162 bp dsRNAs, which can accommodate 1, 2, 3, 4-5 and 11-12 RIG-I molecules, respectively (Figure 3B). The mobility shift assay showed that RIPLET binding requires at least 21 bp dsRNA, *i.e.* two RIG-IΔN molecules in proximity.

RIPLET consists of the N-terminal RING, central coiled coil (CC) and C-terminal PSpry domains (Figure 3B). The RING domain is known to recognize E2 conjugating enzymes, and is essential for both RIG-I ubiquitination and unanchored Ub_n_ synthesis (Figure S3D). RING deletion (ΔRING) had little impact on RIG-I binding (Figure 3B), consistent with the role of RING in E2 engagement rather than substrate binding. Further deletion of the CC domain, however, abolished RIG-I binding (Figure 3C). Since CC is responsible for dimerization of RIPLET (Figure S3E), we examined the importance of dimerization by replacing CC by an orthogonal dimeric protein, GST. The GST-PSpry fusion fully restored RIG-I binding (Figures 3C and S3F), suggesting that PSpry is the RIG-I binding domain, and bivalency is required for high affinity binding (Figure 3D). This is in line with the observation that RIPLET binds RIG-I only in complex with dsRNA that is long enough to accommodate at least two RIG-I molecules (Figure 3B).

We next explored how the bivalent binding mode of RIPLET affects dsRNA length selectivity of RIG-I. We expected that RIPLET would bind and ubiquitinate RIG-I filaments on longer dsRNA more efficiently, because longer filaments would allow many more RIPLET binding configurations (Figure 3D). Comparison of the ubiquitination efficiency, however, showed that RIPLET ubiquitinates RIG-I most efficiently when RIG-I is bound with ~30-40 bp dsRNAs. This ubiquitination is more efficient than when RIG-I is bound with >40 bp dsRNAs (Figure 3E). Although RIG-I filament formation is known to be limited on >~0.5-1 kb dsRNAs (see Figure S3G for the explanation), which could partly explain the decline in Ub conjugation with ~500 bp dsRNA, the maximum ubiquitination at ~30-40 bp was unexpected given the efficient RIG-I filament formation on ~40-~200 bp dsRNA (Binder et al., 2011; Peisley et al., 2013). Interestingly, the signaling activity of RIG-I also showed a bell-shaped curve, but with the peak position shifted to longer dsRNAs (~160 bp) (Figures 3F & S3H). The inefficient ubiquitination of RIG-I on >~40 bp RNAs and its preference for longer dsRNAs in RIG-I signaling led us to question whether RIPLET binds RIG-I differently on >~40 bp dsRNA, perhaps in a manner conducive to RIG-I signaling, but not for ubiquitination. Intriguingly, with longer dsRNAs, we observed two populations of RIG-I:RIPLET complexes in native gels: those migrating into the gel and those remaining in the gel well (Figure S3I).

### RIPLET cross-bridges RIG-I filaments *in vitro*

To examine how RIPLET binds RIG-I filaments on long dsRNAs and whether RIPLET can induce additional higher-order oligomerization of RIG-I, we employed negative stain EM to visually inspect the RIPLET:RIG-I complex. We chose 162 bp and 512 bp dsRNAs, the two longest dsRNAs used in Figure 3. Consistent with previous findings (Peisley et al., 2013), RIG-I formed filaments only upon dsRNA binding (Figures 4A and S4A). Note that dsRNA alone or RIG-I alone are generally invisible by negative stain EM because of inefficient staining and small sizes. This allows clear identification of RIG-I filaments on dsRNA. Comparison of RIG-I filaments before and after RIPLET addition showed that RIPLET induces RIG-I filament bridging (Figure 4A). This was observed with both162 bp and 512 bp dsRNAs. Similar to the intra-filament binding mode, filament bridging required neither the RING domain of RIPLET (Figure 4A) nor the 2CARD domain of RIG-I (Figure 4B). No such aggregate structure was found in the absence of dsRNA or RIG-I (Figure 4B). RIPLET also did not bridge MDA5 filaments (Figure S4B), consistent with the specificity of RIPLET for RIG-I, not MDA5 (Figure 1D & S3B).

**Fig. 4.**
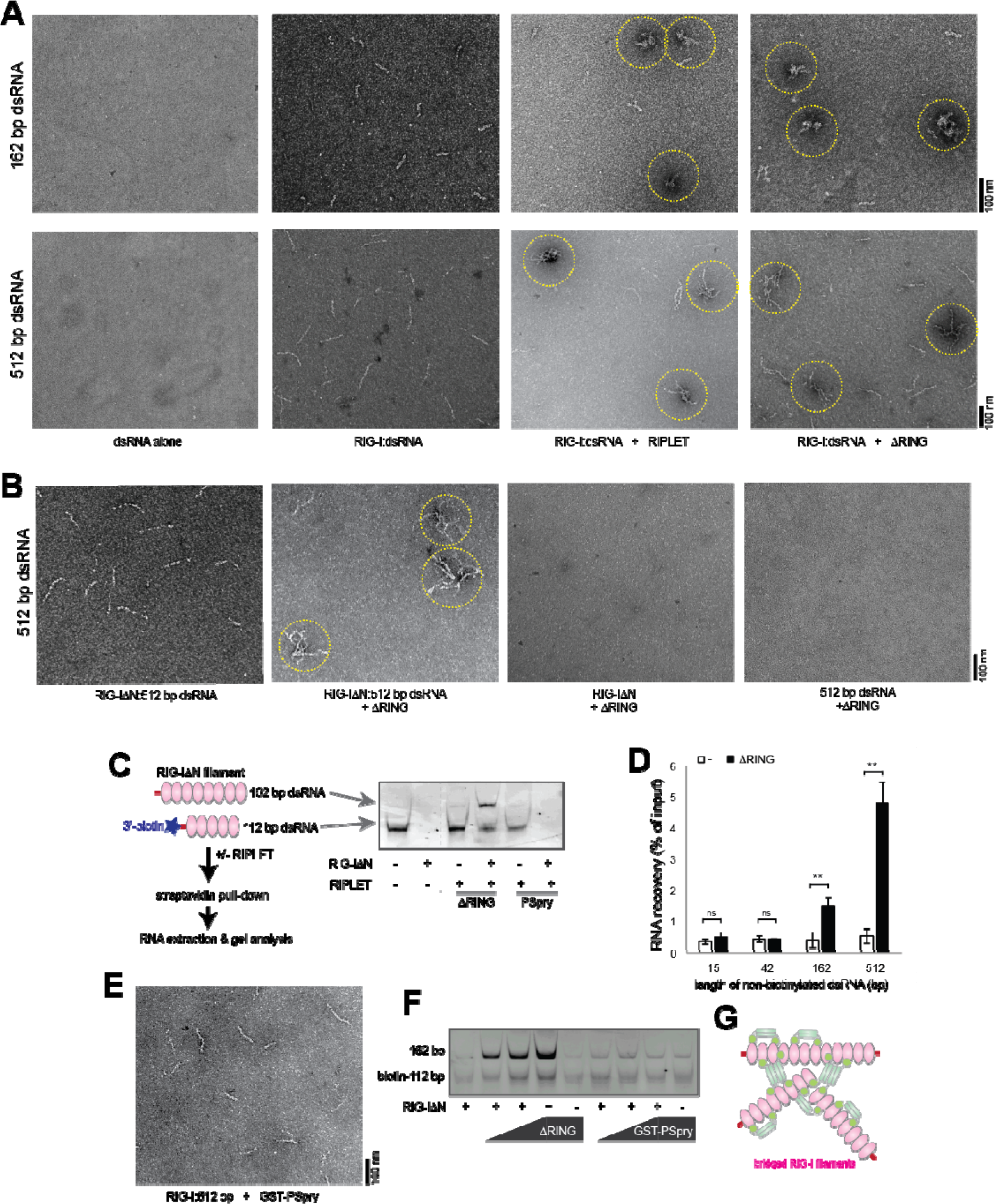
RIPLET cross-bridges RIG-I filaments *in vitro*. A. Representative EM images of RIG-I filaments (0.5 μM) formed on 162 bp (top) or 512 bp (bottom) dsRNA (1 ng/μl), in the presence and absence of RIPLET or ΔRING (0.5 μM). Cross-bridged RIG-I filaments are highlighted with yellow circles. B. Representative EM images of RIG-IΔN filaments (0.5 μM) formed on 512 bp dsRNA (1 ng/μl), in the presence or absence of ΔRING (0.5 μM). Images for ΔRING incubated with RIG-IΔN alone or dsRNA alone were shown on the right as controls. C. Streptavidin pull-down analysis to examine RIPLET-mediated cross-bridging of RIG-I filaments. RIG-IΔN filaments (0.5 μM) were formed on non-biotinylated 162 bp dsRNA (1 ng/μl) and 3’-biotinylated 112 bp dsRNA (1 ng/μl), and the complexes were incubated with ΔRING prior to streptavidin pull-down. The bound RNA was extracted and analyzed by native PAGE (right). D. Recovery rate (mean ± s.d., *n* = 3) of non-biotinylated 21-512 bp dsRNA (1 ng/μl) when incubated with 3’-biotinylated 112 bp dsRNA (1 ng/μl) and RIG-IΔN in the presence and absence of ΔRING, as in (C). * *p* < 0.05, ** *p* < 0.01, ns, not significant (unpaired *t* test, compared with no ΔRING). E. Representative EM image of RIG-I filaments (0.5 μM) formed on 512 bp dsRNA (1 ng/μl), in the presence of GST-PSpry (0.5 μM). F. Streptavidin pull-down analysis using an increasing concentration of ΔRING or GST-PSpry (0.25, 0.5, and 1 μM). The experiment was performed as in (C). G. A model of how RIPLET cross-bridges RIG-I filaments. Weak inter-dimeric interactions between coiled coils (CCs) may allow RIPLET to form multi-valent oligomers (tetramer or larger species) when clustered on a RIG-I filament. Multi-valent RIPLET oligomers would then allow cross-bridging of distinct RIG-I filaments. Note that the two RIPLET binding modes, intra-filament binding (in Figure 3D) and inter-filament bridging, are not mutually exclusive and can co-occur on the same RIG-I filament. All data are representative of three independent experiments.

To independently examine the filament bridging activity, we developed a biotin-RNA pull-down assay, in which the association between biotinylated and non-biotinylated dsRNAs was examined in the presence and absence of RIG-IΔN and RIPLETΔRING (Figure 4C). RIG-IΔN and ΔRING were incubated with a mixture of 3’-biotinylated 112 bp dsRNA and non-biotinylated dsRNA of 162 bp. Biotin-dsRNA and the bound proteins were purified with streptavidin beads, and the level of co-purified, non-biotinylated dsRNA was analyzed by gel. In the absence of RIPLET, co-purification of 162 bp dsRNA and biotin-112 bp dsRNA was minimal and was not improved by the addition of RIG-IΔN (Figure 4C). In fact, RIG-IΔN negatively affected purification of biotin-112 bp dsRNA (Figure 4C), which is likely due to the reduced accessibility of the 3’-conjugated biotin moiety upon RIG-I binding to the dsRNA end. In the presence of ΔRING, however, 162 bp dsRNA was co-purified with biotin-112 bp dsRNA only in the presence of RIG-IΔN (Figure 4C). PSpry did not promote dsRNA co-purification. The requirement of ΔRING and RIG-IΔN for dsRNA co-purification is consistent with the filament bridging activity of RIPLET. Furthermore, comparison of the recovery rate of non-biotinylated 21-512 bp dsRNAs showed that dsRNA co-purification is more efficient with longer dsRNAs (Figure 4D), as would be expected for filament bridging.

To further examine how ΔRING bridges RIG-I filaments, we tested the bridging activity of GST-PSpry, which binds RIG-I filaments as efficiently as ΔRING (Figure S3F). Intriguingly, GST-PSpry did not promote RIG-I filament bridging as measured by both EM and biotin-RNA pull-down assay (Figures 4E and 4F). This result suggests that RIPLET CC is important for RIG-I filament bridging and its role is beyond simple dimerization. To examine whether other types of dimeric CCs can also support RIG-I filament bridging, we used CC adopted from *Arabidopsis thaliana* Mom1 (MCC) or *Saccharomyces cerevisiae* Gal4 (GCC) (Figure S4C). Both PSpry fused to MCC (MCC-PSpry) and GCC (GCC-PSpry) efficiently bound RIG-I filaments (Figure S4D). However, only MCC-PSpry, not GCC-PSpry, induced RIG-I filament bridging (Figure S4D). MCC is a 141 amino acid (aa)-long antiparallel dimer (Nishimura et al., 2012), while GCC is a 16 aa-long parallel CC (Marmorstein et al., 1992). Although the structure of RIPLET CC is not yet available, RIPLET CC is ~64 aa-long (Figure S4E) and is likely an antiparallel dimer. This prediction is based on the fact that its closest homolog is an antiparallel dimeric CC from TRIM25 (Figure S4E), and our Cys-Cys cross-linking data also supports the antiparallel orientation (Figure S4F). Considering that RIPLET CC and MCC, but not GST or GCC, can bridge RIG-I filaments, we speculate that long antiparallel dimeric CCs may have weak affinity for each other. This weak interaction may stabilize when RIPLET is locally concentrated on a RIG-I filament, resulting in the formation of multivalent RIPLET oligomers and bridging of RIG-I filaments (Figure 4G).

### RIPLET cross-bridges RIG-I filaments *in cellulo* independent of stress granules

To further examine whether RIPLET-mediated bridging of RIG-I filaments indeed occurs in cells, we performed a similar biotin-RNA pull-down analysis in WT and RIPLET^−/−^ 293T cells. Upon transfection with biotin-112 bp dsRNA and non-biotinylated 162 bp dsRNA, cells were lysed and subjected to streptavidin pull-down. The level of 162 bp dsRNA co-purified with biotinylated RNA was measured by RT-qPCR. The result showed that 162 bp dsRNA was co-purified in a manner dependent on biotinylated 112 bp dsRNA (Figure 5A). More importantly, co-purification of 112 bp and 162 bp dsRNA was impaired by RIPLET knock-out (Figure 5A) and was restored by ΔRING complementation (Figure 5B). Similarly, non-biotinylated 512 bp dsRNA was co-purified with biotinylated 112 bp in an RIPLET-dependent manner (Figures 5A & 5B). These data suggest that RIPLET bridges RIG-I filaments in cells and at the endogenous level of RIG-I and RIPLET.

**Fig. 5.**
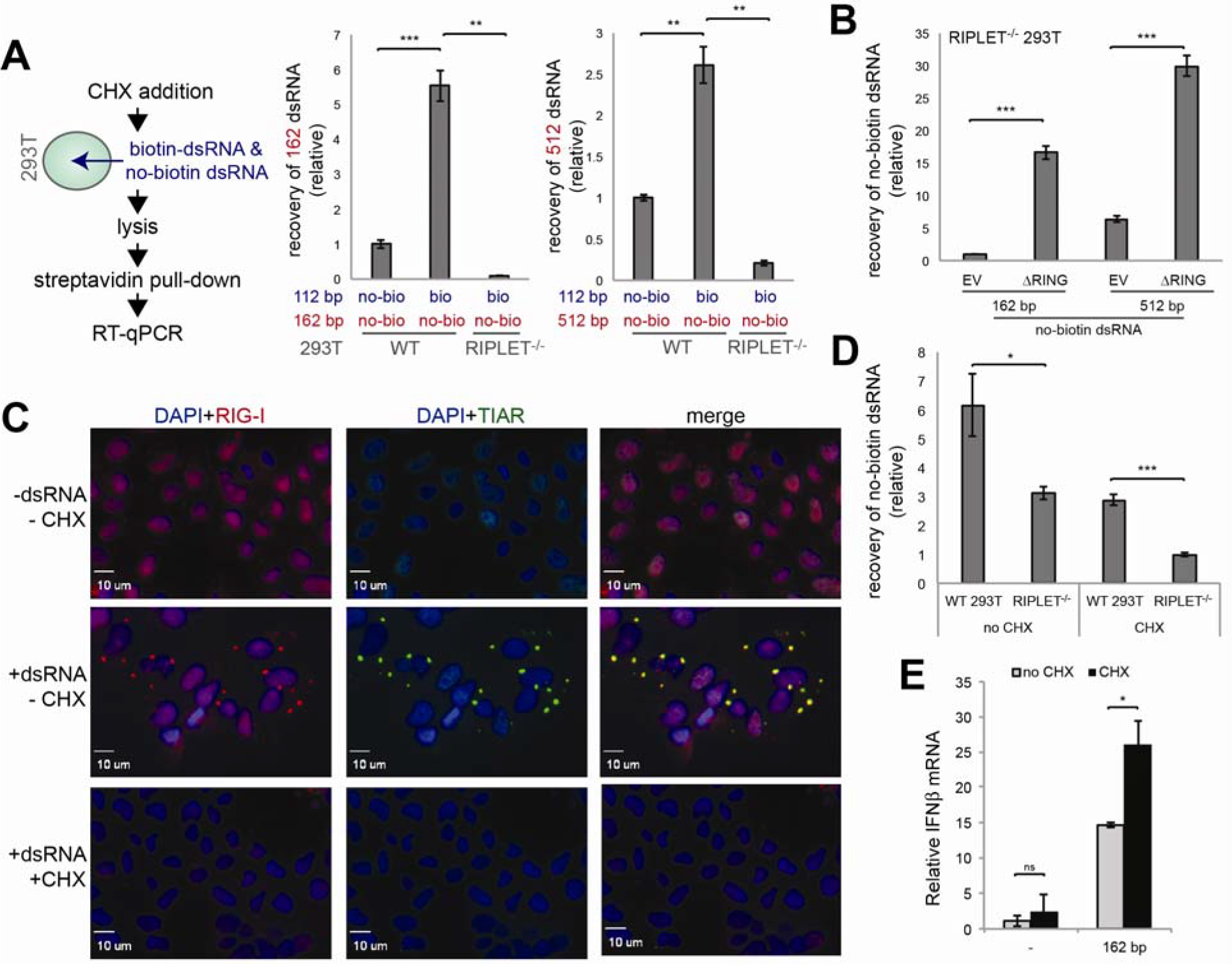
RIPLET cross-bridges RIG-I filaments *in cellulo* independent of stress granules. A. *In cellulo* strepavidin pull-down analysis. WT and RIPLET^−/−^ 293T cells were treated with cyclohexamide (CHX, 0.1 mg/ml) an hour before transfection with non-biotinylated dsRNA (either 162 bp or 512 bp, 1 μg in a 6 well plate) and 3’-biotinylated 112 b dsRNA (1 μg). Cells were lysed 4 hrs post RNA transfection and were subjected to streptavidin pull-down as described in STAR methods. Co-purification of non-biotinylated 162 bp or 512 bp dsRNA was measured by RT-qPCR. No-bio and bio refer to non-biotinylated dsRNA and biotinylated dsRNA, respectively. B. *In cellulo* strepavidin pull-down analysis using 162/512 bp dsRNA (1 μg) and 3’-biotinylated 112 bp dsRNA (1 μg). The ΔRING construct (or empty vector, EV) was transfected into RIPLET^−/−^ 293T cells 24 hr before dsRNA transfection. Experiments were performed as in (A). C. Immunofluorescence analysis of endogenous RIG-I and SGs (marked by αTIAR staining) in 293T cells in the presence and absence of 162 bp dsRNA (0.5 μg) and CHX. Cells were treated with CHX (0.1 mg/ml) an hour before transfection with dsRNA (0.2 μg/ml), and fixed for imaging 2 hr post dsRNA transfection. D. *In cellulo* strepavidin pull-down analysis with and without CHX (0.1 mg/ml). 162 bp dsRNA was used as no-biotin dsRNA. Experiments were performed as in (A). E. The effect of CHX (0.1 mg/ml) on RIG-I signaling in response to 162 bp dsRNA. All data are presented as mean ± s.d. (*n* = 3 for A-D and *n* = 2 for E) and are representative of three independent experiments. * *p* < 0.05, ** *p* < 0.01, \*\*\**p* < 0.001, ns, not significant (unpaired *t* test).

Note that the observed co-purification of dsRNAs is independent of stress granules (SGs). SGs are cytosolic phase separation structures formed by RNA and RNA-binding proteins including RIG-I (Oh et al., 2016; Onomoto et al., 2013), and are known to form upon dsRNA stimulation (Figure 5C). Although a previous study suggested a role of RIPLET in SG formation (Oshiumi et al., 2013), our analysis indicates an effect of RIPLET on the kinetics of RIG-I SG formation, rather than on its steady state level (Figures S5A-B). To avoid any potential confounding effect of SGs on RIG-I filament bridging, we used cyclohexamide (CHX), which is known to inhibit SG formation (Figure 5C). In the absence of CHX, we still observed RIPLET-dependent RNA bridging, but the use of CHX significantly reduced the background RNA bridging (Figure 5D). Note that Figures 5A-B were performed in the presence of CHX (see the flow chart in Figure 5A). The observed co-purification of two RNAs, even in the presence of CHX, suggests that RIPLET-mediated filament bridging occurs independent of SGs. CHX also did not negatively affect RIG-I signaling (Figure 5E), suggesting that SG localization of RIG-I, which is distinct from RIPLET-mediated filament bridging, is not important for antiviral signaling.

### RIG-I signaling is enhanced by RIPLET-mediated filament cross-bridging

We next asked whether filament bridging contributes to RIG-I signaling. On one hand, filament bridging would cluster RIG-I molecules, which could promote 2CARD tetramerization and thus, MAVS activation, with or without K63-Ub_n_. On the other hand, SG assembly, which also clusters RIG-I in a RIPLET-independent manner, had no impact on RIG-I signaling (Figure 5E). To test the potential role of RIPLET-mediated filament bridging, we first employed the *in vitro* MAVS polymerization assay as in Figures 2D & 2E. Because this assay uses only the purified proteins and RNA, it allows one to directly measure the impact of filament bridging on RIG-I signaling in the absence of other cellular components. Since K63-Ub_n_ can be introduced *in trans*, this assay can also test the effect of RIG-I filament bridging, both in the presence and absence of K63-Ub_n_. While K63-Ub_n_ alone stimulated RIG-I’s ability to induce MAVS filamentation, ΔRING significantly improved the MAVS stimulatory activity of RIG-I (Figure 6A). Furthermore, even in the absence of K63-Ub_n_, ΔRING stimulated MAVS filamentation (Figure 6B), albeit not as efficiently as in the presence of K63-Ub_n_. GST-PSpry, by contrast, had no such stimulatory effect (Figure 6B), consistent with the role of RIG-I filament bridging in MAVS activation. The significantly more efficient MAVS filamentation in the presence of both K63-Ub_n_ and ΔRING (Figure 6A), compared to either alone, suggests that the two are synergistic, rather than having an additive effect on RIG-I signaling.

**Fig. 6.**
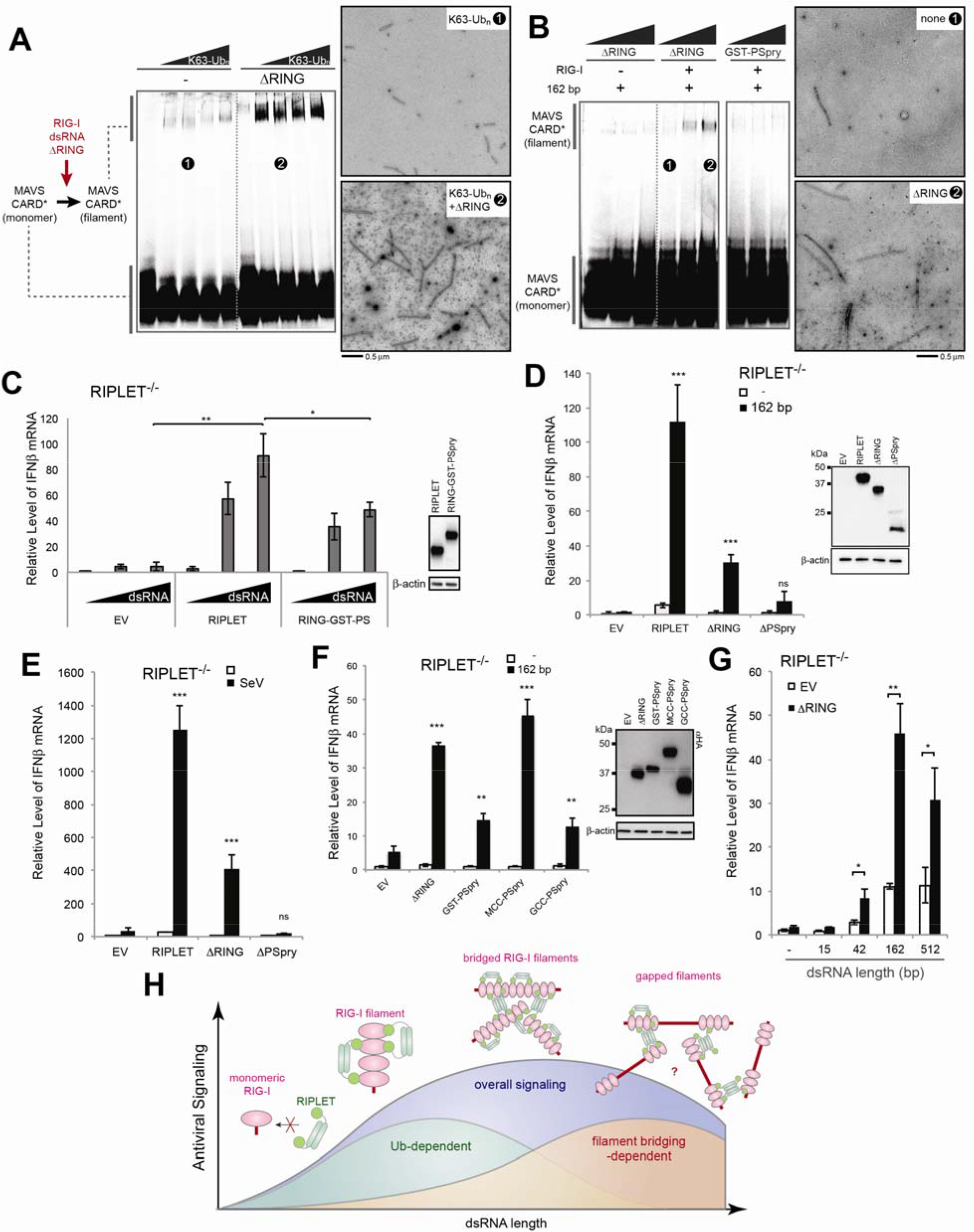
RIPLET-mediated RIG-I aggregation enhances antiviral signaling, in both ubiquitin-dependent and -independent manners. A. MAVS polymerization assay to examine the effect of ΔRING (12.5 μM) in the presence of an increasing concentration of K63-Ub_n_ (equivalent to 0-7 μM monomeric Ub). Right: representative EM images of MAVS CARD filaments nucleated with RIG-I filaments without ΔRING (*) or with ΔRING (**), all in the presence of K63-Ub_n_ (equivalent to 1.4 μM monomeric Ub). B. MAVS polymerization assay to examine the effect of ΔRING in the absence of K63-Ub_n_. The RIG-I:dsRNA complex (1 μM RIG-I and 6 ng/μl 162 bp dsRNA) was incubated with an increasing concentration of ΔRING or GST-PSpry (0, 5, 12.5 μM), and mixed with purified monomeric MAVS CARD (10 μM). Polymerization of MAVS CARD was measured by native PAGE and EM. Right: representative images for MAVS CARD filaments nucleated with RIG-I filaments without ΔRING (*) or with ΔRING (**). C. Comparison between full-length RIPLET and RING fused to GST-PSpry (RING-GST-PSpry) (20 ng each) in the RIG-I stimulatory activity. Relative level of antiviral signaling was measured upon stimulation with 162 bp dsRNA (0.2 μg) in RIPLET^−/−^ 293T. D-E. Relative level of RIG-I signaling upon stimulation with 162 bp dsRNA (0.2 μg) (D) or SeV (100 HA units/ml) (E) in RIPLET^−/−^ 293T cells complemented with empty vector (EV), RIPLET, ΔRING or ΔPSpry (20 ng). The level of antiviral signaling was measured by IFNβ mRNA. All cells were transiently transfected with the RIG-I expression plasmid (10 ng). See Figure S6 for the effect of ΔRING on endogenous RIG-I. F. Effect of ΔRING and CC-swap mutants on RIG-I signaling in RIPLET^−/−^ 293T upon stimulation with 162 bp dsRNA. Experiments were performed as in (D). G. dsRNA length-dependence of the effect of ΔRING on RIG-I signaling. Experiments were performed as in (D). H. A model of how RIPLET promotes RIG-I signaling using both Ub-dependent and – independent mechanisms. RIG-I first binds a dsRNA end as a monomer, but on long dsRNAs, it forms filamentous oligomers. RIPLET selectively binds and ubiquitinates filamentous RIG-I because bivalent binding is required for their high affinity interactions. On ~40-~500 bp RNAs, RIPLET can also cross-bridge RIG-I filaments, leading to the higher-order receptor clustering and amplification of antiviral signaling. On much longer (>~1 kb) dsRNA, however, RIG-I inefficiently forms filaments (due to the relative dilution of 5’ppp-harboring end), which consequently limits RIPLET binding, RIG-I clustering and antiviral signaling. Our data suggest that the effect of RIG-I clustering synergizes with Ub-mediated oligomerization of 2CARD (not depicted in the cartoon) to amplify RIG-I signaling on the basis of the duplex RNA length. Because they act synergistically, not additively, the relative contributions of Ub-dependent and filament bridging-dependent mechanisms are difficult to quantitate and are depicted with arbitrary amplitudes. All data are presented as mean ± s.d. (*n* = 3-4) and are representative of three independent experiments. * *p* < 0.05, ** *p* < 0.01, \*\*\**p* < 0.001, ns, not significant (unpaired *t* test). Unless indicated otherwise, *p* values were calculated in comparison to EV.

To further test the effect of RIG-I filament bridging in cells, we compared the RIG-I-stimulatory activity of full-length RIPLET and RING fused to GST-PSpry (RING-GST-PSpry) in RIPLET^−/−^ 293T cells. RIPLET can bridge RIG-I filaments, while GST-PSpry cannot (Figures S3F & 4E). The result shows that RIPLET is a more potent stimulator of RIG-I than is RING-GST-PSpry (Figure 6C), consistent with the notion that RIG-I filament bridging stimulates RIG-I signaling. Furthermore, ΔRING, which can bridge RIG-I filaments but cannot ubiquitinate RIG-I (Figures 4A & S3D), also enhanced RIG-I signaling in response to dsRNA and viral infection (Figures 6D & 6E), albeit not as efficiently as full-length RIPLET. The RIG-I-stimulatory effect of ΔRING was observed with both transiently expressed ΔRING (Figures 6D & 6E) and stably reconstituted ΔRING (Figure S6A).

To further examine whether the positive effect of ΔRING on RIG-I signaling is mediated by inter-filament bridging, we compared the RIG-I-stimulatory activity of ΔRING and its CC-swap mutants, which showed varying degrees of filament bridging (Figures 4F & S4D). Parallel to their abilities to bridge RIG-I filaments, ΔRING and MCC-PSpry robustly stimulated RIG-I signaling, but GST-PSpry and GCC-PSpry had only a moderate effect (Figure 6F). Furthermore, comparison among dsRNAs of various lengths suggests that ΔRING-mediated RIG-I signaling is more pronounced with longer RNA (Figure 6G), consistent with the length-dependence of filament bridging. While the exact pattern of dsRNA length dependence in Figure 6G differs from that of *in vitro* filament bridging in Figure 4D, this is likely because the latter is the measure of RNA bridging, not the level of RIG-I clustering and its impact on signaling. Altogether, these observations collectively support the notion that RIPLET uses the filament bridging activity to amplify RIG-I signaling and this activity synergizes with the Ub-dependent RIG-I stimulatory activity (Figure 6H).

## DISCUSSION

According to the current notion in the field of RIG-I signaling, RIG-I is activated by dsRNA binding and subsequently by TRIM25-mediated ubiquitination. This was largely based on the studies of isolated 2CARD, the signaling domain of RIG-I. RIPLET was later identified as a required protein for RIG-I signaling, but was shown to lack the stimulatory effect on isolated 2CARD (Gao et al., 2009; Oshiumi et al., 2009; Oshiumi et al., 2010). Based on the signaling activity of isolated 2CARD, which partially depends on TRIM25 but not on RIPLET, RIPLET was proposed to function upstream of TRIM25 in releasing the auto-repression of 2CARD (Oshiumi et al., 2013). We here showed that RIPLET, but not TRIM25, is required for full-length RIG-I signaling (Figures 1 and S1), and that the mode of action of RIPLET differs from what was previously proposed. While we did observe a moderate (~2-fold) stimulatory effect of TRIM25 on the signaling activity of isolated RIG-I 2CARD, as previously reported (Gack et al., 2007) (Figures S1H & S1I), knocking-out TRIM25 had little negative impact on full-length RIG-I signaling (Figures 1 and S1). This was demonstrated using multiple, independent knock-out cell lines in three different cell types (293T, MEF, A549) with five different RIG-I stimuli (dsRNA, SeV, VSV, RVFV and FluA). The lack of impact of TRIM25 on RIG-I function was surprising given the widely held view of its essential role in RIG-I signaling. Considering the growing list of RIG-I-independent functions of TRIM25 in antiviral immunity–in part mediated by its RNA binding activity (Choudhury et al., 2014; Manokaran et al., 2015; Meyerson et al., 2017) and its interaction with ZAP (Li et al., 2017; Zheng et al., 2017)–the precise nature of the observed effect of TRIM25 on RIG-I 2CARD requires more detailed investigation.

Using a combination of biochemical analyses, we also showed that RIPLET is sufficient to conjugate full-length RIG-I with K63-Ub_n_ and to stimulate MAVS filamentation (Figure 2). TRIM25 does not ubiquitinate full-length RIG-I as robustly as RIPLET (Figure 2A). While previous studies, including our own (Peisley et al., 2014), described ubiquitination of isolated 2CARD by TRIM25, this required significantly higher concentrations of E1, E2, E3 enzymes and substrates. More importantly, ubiquitination of RIG-I by TRIM25 is not stimulated by dsRNA (Figure 2A), further raising a question on the role of TRIM25 in RIG-I ubiquitination. By contrast, RIPLET binds and ubiquitinates RIG-I only upon dsRNA binding (Figures 2 and 3), the mode of action consistent with the cellular activation condition for RIG-I signaling. Of note, RIPLET also promotes unanchored Ub_n_ synthesis, but this activity is suppressed upon binding to RIG-I (Figure 2G). Additionally, the PSpry domain of RIPLET, which is dispensable for unanchored Ub_n_ synthesis, is required for conjugating Ub_n_ to RIG-I and for activating RIG-I signaling (Figures 2H & 2I). These results further suggest the role of anchored K63-Ub_n_ chains, not unanchored ones, in RIG-I signaling.

Our biochemical analyses of RIPLET revealed a novel mechanism by which RIPLET recognizes RIG-I in an oligomeric state-specific manner, and further induces higher order receptor “clustering”. RIPLET binds the core RIG-I filaments formed by the helicase domain and CTD, and ubiquitinates multiple sites on the RIG-I protein, including 2CARD. More detailed analysis of the interactions between RIPLET and RIG-I showed that RIPLET utilizes its dimeric structure and the bivalent binding mode to preferentially recognize and ubiquitinate filamentous oligomers of RIG-I on >20 bp dsRNA. On longer dsRNA, RIPLET can also cross-bridge RIG-I filaments in an Ub-independent manner. This bridging further promotes the signaling activity of RIG-I, likely by inducing high local concentrations of the receptor, thereby facilitating 2CARD tetramerization and, hence, nucleation of MAVS filaments. The combination of the intra-filament binding and inter-filament bridging modes enables RIG-I to signal more efficiently on longer dsRNA (Figure 6H). In other words, RIPLET utilizes both Ub-dependent and – independent mechanisms to amplify RIG-I signaling on the basis of dsRNA length.

Based on the role of RIPLET in both RNA-discrimination and receptor oligomerization, we propose that RIPLET functions more like a co-receptor, rather than a downstream accessory protein. Our findings on RIPLET provide a previously unrecognized mechanism for how the innate immune system utilizes an E3 ligase to measure foreign nucleic acid length, a criterion commonly used by DNA and RNA sensors for self vs. non-self discrimination (Andreeva et al., 2017; Morrone et al., 2014; Peisley et al., 2012; Zhou et al., 2018). Additionally, RIPLET-mediated clustering of RIG-I provides a new perspective on receptor clustering, a phenomenon studied primarily in the context of cell-surface receptors (Hartman and Groves, 2011). It is yet unclear exactly how RIPLET specifically recognizes RIG-I filaments and discriminates against MDA5 filaments. Considering that a viral protein can distinguish between RIG-I and MDA5 using a single amino acid difference (Motz et al., 2013; Rodriguez and Horvath, 2013), one could speculate an analogous, sequence-dependent recognition mechanism for RIPLET. Future research is required to understand how RIPLET specifically recognizes RIG-I, and which E3 ligase recognizes MDA5 and how. Given the importance of the coiled-coil dimeric architecture in RIPLET functions and the commonality of this architecture in many E3 ligases (e.g. TRIM family members), our findings on RIPLET may be widely applicable to other E3 ligases recognizing MDA5 and beyond.

## Acknowledgement

We thank Sandra Wüst for her assistance, Drs. Gack and Oshiumi for sharing TRIM25^−/−^ and RIPLET^−/−^ MEFs, respectively, and Dr. O.Pornillos for sharing the TRIM25 construct and protein purification protocol. This work was supported by NSF fellowship (C.C.), Cancer Research Institute fellowship (S.A.), German Academic Exchange Service PROMOS program (A.X.), the German Research Foundation (DFG TRR179, TP11) (M.B.), German Federal Ministry of Education and Research (ERASysApp, 031A602C) (M.B.), Burroughs Wellcome Fund and NIH grants (R01AI106912, R01AI111784 and R21AI130791) (S.H.).

## Author contributions

CC, SA, AX, JW, JWP, MB and SH designed experiments and analyzed data. CC, SA, AX, JW, SP and JWP performed experiments. SWO and TF provided αRIG-I antibody. FH and MB provided knock-out cell lines and FH provided αRIPLET antibody.

---

**Supplementary Fig. 1.**
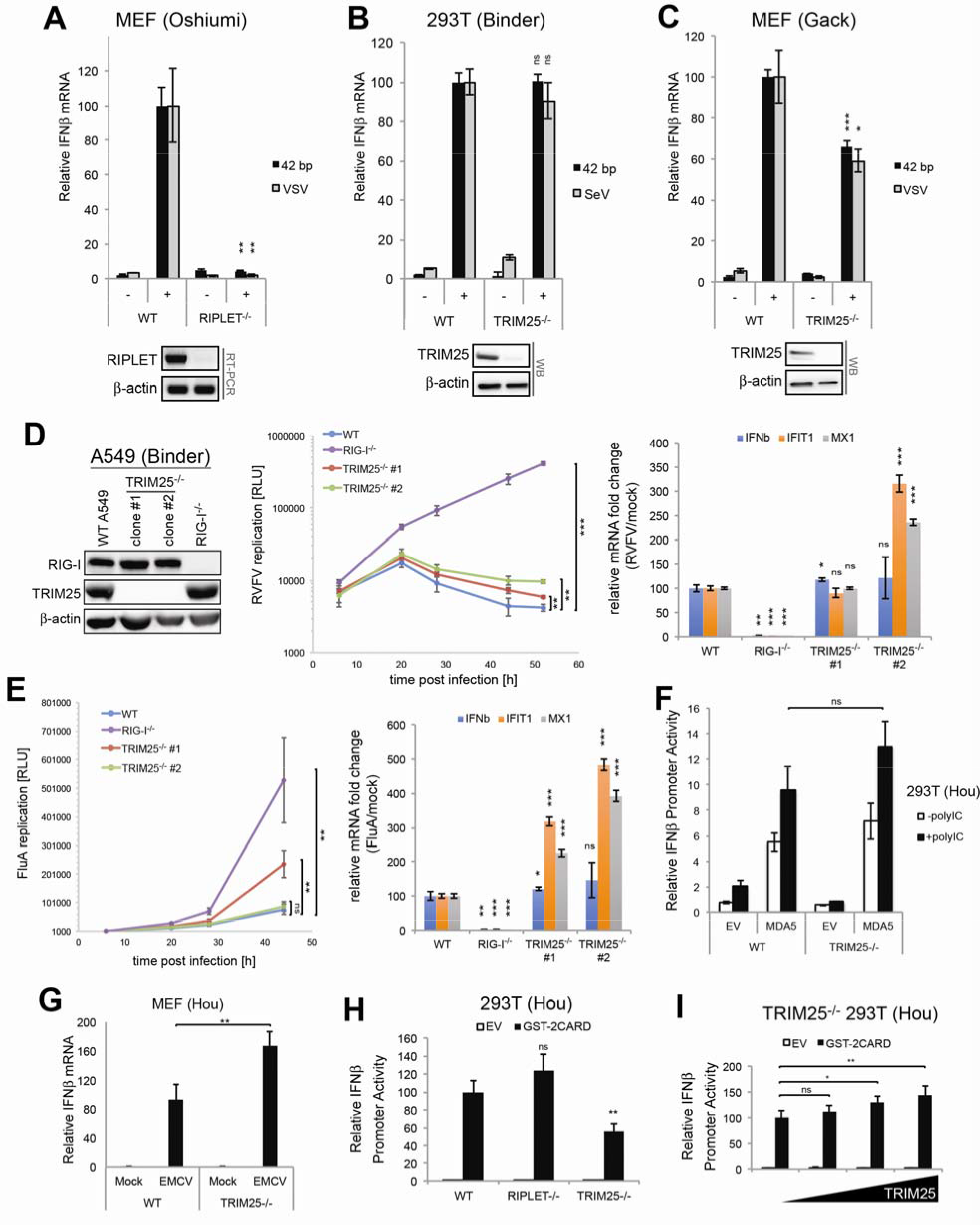
RIPLET, not TRIM25, is required for RIG-I signal activation. A-C. The effect of RIPLET or TRIM25 deletion on antiviral signaling. MEF (WT and RIPLET^−/−^), 293T cells (WT and TRIM25^−/−^) and MEF (WT and TRIM25^−/−^) were obtained from Oshiumi (Oshiumi et al., 2010), Binder (see STAR Methods) and Gack (Gack et al., 2007) laboratories, respectively. Cells were stimulated with 42 bp dsRNA with 5’ppp (0.2 μg), SeV (100 HA units/ml) or VSV (MOI=1), and the antiviral signaling activity was measured by the IFNβ mRNA level (using GAPDH as an internal control), and normalized against the WT values. D-E. Virus replication and antiviral signaling in WT (non-targeting CRISPR control), RIG-I^−/−^ and TRIM25^−/−^ A549 cells. Luciferase-expressing RVFV (RVFVΔNSs_Rluc, MOI=0.05) (D) and FluA (FluA_GLuc, MOI=0.001) (E) were used (see STAR methods). The level of viral replication was measured by the luciferase activity. The level of antiviral signaling was measured by the mRNA levels of IFNβ IFIT1 and Mx1 (using GAPDH as an internal control) at 20 hr post-infection for RVFV and 40 hr post-infection for FluA. All values were normalized against the WT values. Unlike other experiments in this manuscript, these experiments were performed in the Binder laboratory for an independent evaluation of the role of TRIM25 in RIG-I signaling.
a. Relative signaling activity of MDA5 in WT vs. TRIM25^−/−^ 293T cells (from the Hou lab), as measured by the IFNβ promoter reporter assay. Cells were transiently transfected with MDA5 expression vector (5 ng) and were stimulated with polyIC (0.5 μg) as in Figure 1D.
b. Relative signaling activity of endogenous MDA5 in WT vs. TRIM25^−/−^ MEF cells (from the Hou lab), as measured by the IFNβ mRNA level. Cells were stimulated with encephalomyocarditis virus (EMCV, MOI= 0.1), which is known to be recognized by MDA5, not RIG-I.
c. Relative signaling activity of RIG-I 2CARD fused to GST (GST-2CARD) (50 ng) in 293T cells (WT, RIPLET^−/−^ or TRIM25^−/−^, from the Hou lab (Shi et al., 2017)), as measured by the IFNβ promoter reporter assay. Note that antiviral signaling by GST-2CARD occurs in a RIPLET-independent manner because, as shown in Figure 3A, RIPLET does not recognize isolated 2CARD.
d. Relative signaling activity of GST-2CARD in TRIM25^−/−^ 293T cells (from the Hou lab) with and without ectopically expressed TRIM25. An increasing amount of TRIM25 expression construct (0, 2, 10, 50 ng) was co-transfected with the GST-2CARD expression construct or empty vector (EV) (50 ng). All data are presented as mean ± s.d. (*n* = 3-4) and are representative of three independent experiments. * *p* < 0.05, ** *p* < 0.01, \*\*\**p* < 0.001, ns, not significant (unpaired *t* test, compared with WT values).

**Supplementary Fig. 2.**
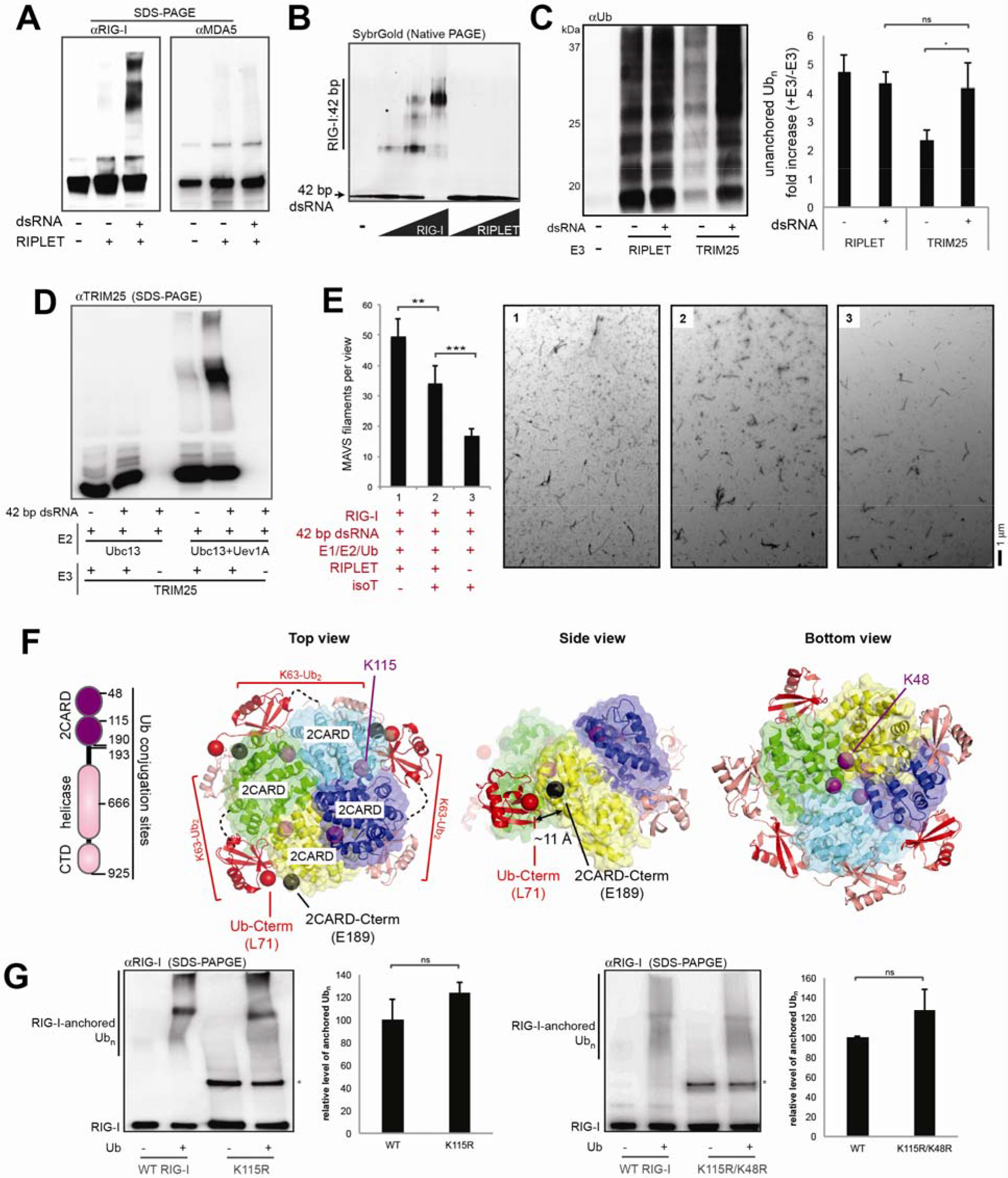
RIPLET, not TRIM25, ubiquitinates RIG-I in a dsRNA-dependent manner. a. *In vitro* ubiquitination of RIG-I and MDA5 by RIPLET. RIG-I or MDA5 (0.5 μM) was incubated with RIPLET (0.25 μM) in the presence or absence of their preferred dsRNA substrates (42 bp for RIG-I and 512 bp for MDA5, 1 ng/μl). Reactions were performed as in Figure 2A, and analyzed by anti-RIG-I or anti-MDA5 blot.
b. RNA binding activity of RIG-I and RIPLET, as measured by native gel shift assay. An increasing concentration of RIG-I or RIPLET (0.25, 0.5, 1 μM) was incubated with 42 bp dsRNA and analyzed by native PAGE. Gel image was obtained using fluorescence of RNA stained with SybrGold.
c. Unanchored ubiquitin synthesis by RIPLET and TRIM25 in the presence or absence of 42 bp dsRNA. All reactions were performed as in Figure 2A, but RIG-I was omitted in the reaction because RIPLET generates unanchored Ub chains more efficiently in the absence of RIG-I (see Figure 2G). Unanchored Ub chains were analyzed by anti-Ub blot as in Figure 2G. Right: quantitation of unanchored Ub chains (mean ± s.d., *n* = 3). * *p* < 0.05, ** *p* < 0.01, \*\*\**p* < 0.001, ns, not significant (unpaired *t* test).
d. Analysis of TRIM25 auto-ubiquitination. Samples in Figure 2A were re-analyzed by anti-TRIM25 blot.
e. MAVS polymerization assay. RIG-I was subjected to the ubiquitination reaction as in Figure 2A, and subsequently treated with isoT prior to mixing with MAVS CARD (10 μM). MAVS CARD polymerization reaction was carried out as in Figure 2D, and analyzed by negative stain EM as in Figure 2E. Briefly, six random images were collected at 6,800x magnification (right), and average number (± s.d.) of MAVS filaments per image was plotted (left). The result suggests that RIG-I-induced MAVS stimulation is most efficient after the RIPLET reaction before isoT treatment (sample 1). IsoT somewhat reduces the MAVS stimulatory activity of RIG-I (sample 2), but is still more efficient than RIG-I without RIPLET (sample 3). The moderate effect of isoT could be due to partial degradation of anchored Ub chains and/or complete degradation of unanchored Ub chains (see Figure 2F).
f. Mass-spectrometry analysis of Ub conjugation sites in RIG-I. Ubiquitination reaction was performed as in Figure 2A in the presence of 42 bp dsRNA. Ub conjugation was detected at multiple sites within RIG-I (left). Among those, K190 and K193 reside within the flexible C-terminal region of 2CARD (residues ~190-200) located in close proximity to the Ub binding site. See the top and side views of the crystal structure PDB:4NQK (Peisley et al., 2014). This observation supports the notion that ubiquitination at K190 and K193 promotes Ub:2CARD interaction and thus the 2CARD tetramer formation and antiviral signaling. In addition to K190 and K193, mass-spectrometry analysis also identified K48 and K115 in 2CARD to be ubiquitinated by RIPLET. K48 and K115 are near and at the 2CARD tetramerization interface, respectively (see top and bottom views). Accordingly, ubiquitination at these sites may interfere with 2CARD tetramerization. However, quantitative analysis in (G) suggests that these two sites are ubiquitinated at low frequencies. Black spheres indicate the most C-terminal residues in individual 2CARD subunits (residue ~190). Red spheres indicate the most C-terminal residues of individual Ub molecules.
g. Contribution of K48 and K115 residues on RIG-I Ub conjugation. WT RIG-I, K115R and K48R/K115R were ubiquitinated as in Figure 2A and analyzed by αRIG-I WB. The result shows that K115R and K48R/K115R mutations have little impact on the overall level of RIG-I ubiquitination, suggesting that K48 and K115 are ubiquitinated at low frequencies. Data are represented as mean ± s.d., *n* = 3. ns, not significant *p* > 0.05 (unpaired *t* test). * likely the uncleaved NusA-RIG-I fusion protein, from which RIG-I is purified.

**Supplementary Fig. 3.**
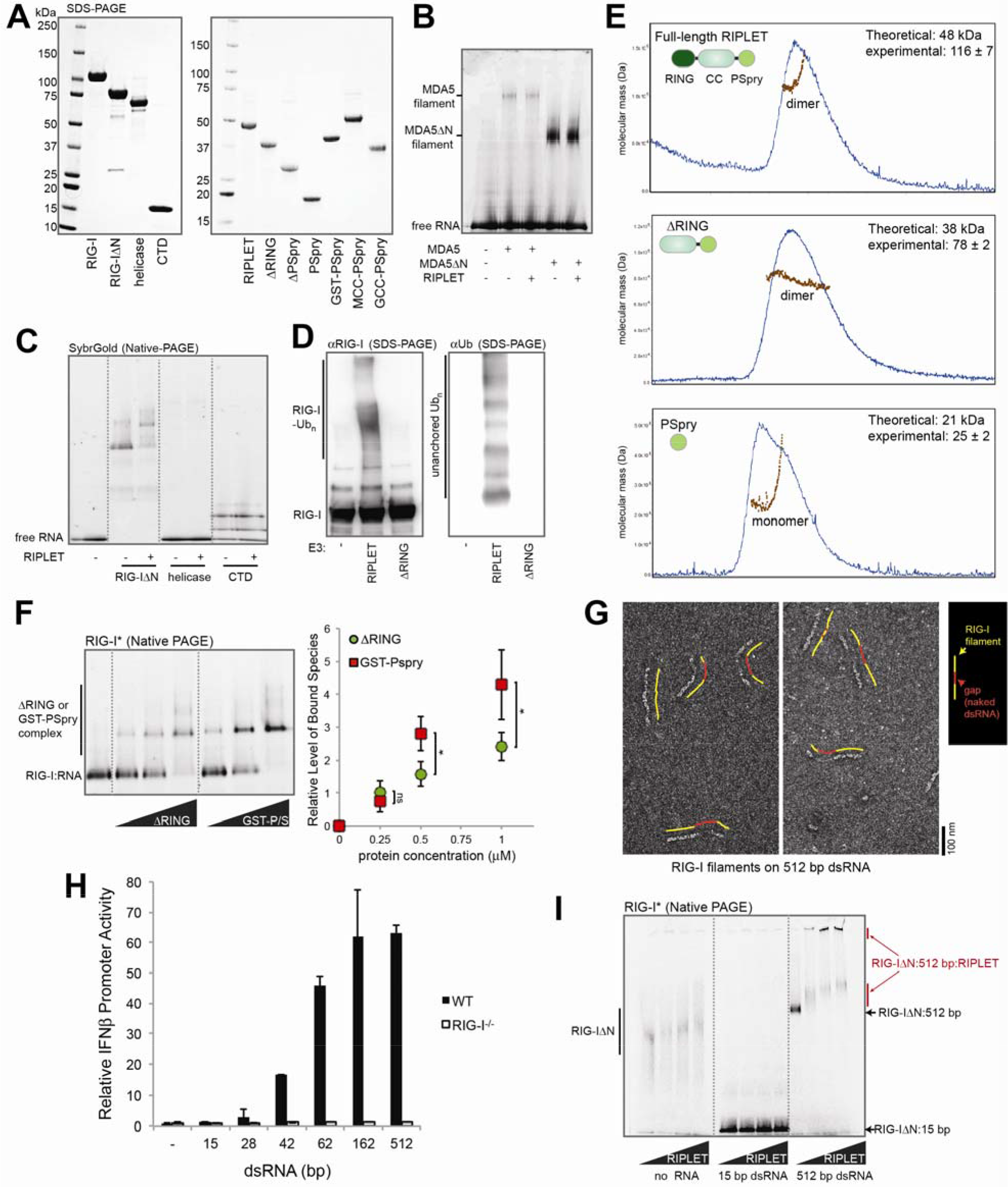
RIPLET utilizes bivalency to selectively recognize RIG-I on pre-oligomerized dsRNA. a. SDS-PAGE analysis of purified RIG-I and RIPLET variants used in this study.
b. Native gel shift assay of MDA5 with RIPLET. 512 bp dsRNA (1 ng/μl) was incubated with MDA5 or MDA5ΔN (0.5 μM), in the presence or absence of RIPLET (0.5 μM). 512 bp dsRNA was visualized using fluorescence of the SybrGold stain.
c. Native gel shift assay of RIG-I variants with RIPLET. 42 bp dsRNA (1 ng/μl) was incubated with RIG-IΔN, the helicase domain or CTD (0.5 μM), in the presence or absence of RIPLET (0.5 μM). 42 bp dsRNA was visualized using fluorescence of the SybrGold stain. The helicase domain does not stably associate with dsRNA, and therefore does not bind RIPLET. Isolated CTD binds dsRNA, but does not bind RIPLET.
d. *In vitro* Ub chain synthesis by RIPLET and ΔRING. Left: Ub conjugation reaction was performed in the presence of 42 bp dsRNA as in Figure 2A, and was analyzed by anti-RIG-I blot. Right: unanchored Ub synthesis reaction was performed without RIG-I or 42 bp dsRNA, because the presence of RIG-I and dsRNA suppresses unanchored Ub synthesis (Figure 2G). Unanchored Ub chains were analyzed by anti-Ub blot. As in Figure 2F, the two blots cover different molecular weight ranges; the anti-RIG-I blot is for >~100 kDa, whereas the anti-Ub blot is for <~50 kDa.
e. Multi-angle light scattering (MALS) traces of full-length RIPLET, ΔRING and PSpry. Theoretical and experimental molecular weights are shown on the upper right corner. Only PSpry exists as a monomer, while full-length RIPLET and ΔRING are constitutive dimers. Blue and brown traces indicate OD_280_ and estimated molecular weight, respectively.
f. Native gel shift assay to compare the RIG-I binding efficiency of ΔRING and GST-PSpry. The assay was performed as in (B). Right: quantitation of the bound species (mean ± s.d., *n* = 3). * *p* < 0.05, ** *p* < 0.01, \*\*\**p* < 0.001, ns, not significant (unpaired *t* test).
g. Representative negative stain electron micrographs of RIG-I filaments on 512 bp dsRNAs. Filaments were formed by incubating dsRNA (1 ng/μl) and RIG-I (0.5 μM) with ATP (2 mM) at 37°C for 10 min, prior to the grid preparation. Note that naked dsRNA is much thinner, allowing unambiguous distinction between RIG-I-coated regions (indicated by yellow lines) and naked regions (red lines) of dsRNA. As evidenced by gaps in the RIG-I filaments, filament assembly of RIG-I on >~0.5-1 kb dsRNA is limited. This is because the filament assembles from a dsRNA end and its propagation to the dsRNA interior is relatively inefficient (Peisley et al., 2013).
h. Relative level of antiviral signaling in response to 15-512 bp dsRNA (0.2 μg for all RNAs) in 293T cells (WT or RIG-I^−/−^), as measured by the IFNβ promoter reporter assay (mean ± s.d., *n* = 2).
i. Native gel shift assay to monitor the interaction between RIG-I and RIPLET. Fluorescently labeled RIG-IΔN (300 nM) was incubated with RIPLET (0, 150, 300, 600 nM) in the presence or absence of 15 bp or 512 bp dsRNA (2.5 ng/μl), and the complex was analyzed by native gel using RIG-I fluorescence. Note that there are two populations of the RIG-I:512 bp:RIPLET complex, one migrating into the gel and the other remaining in the gel well. All data are representative of three independent experiments.

**Supplementary Fig. 4.**
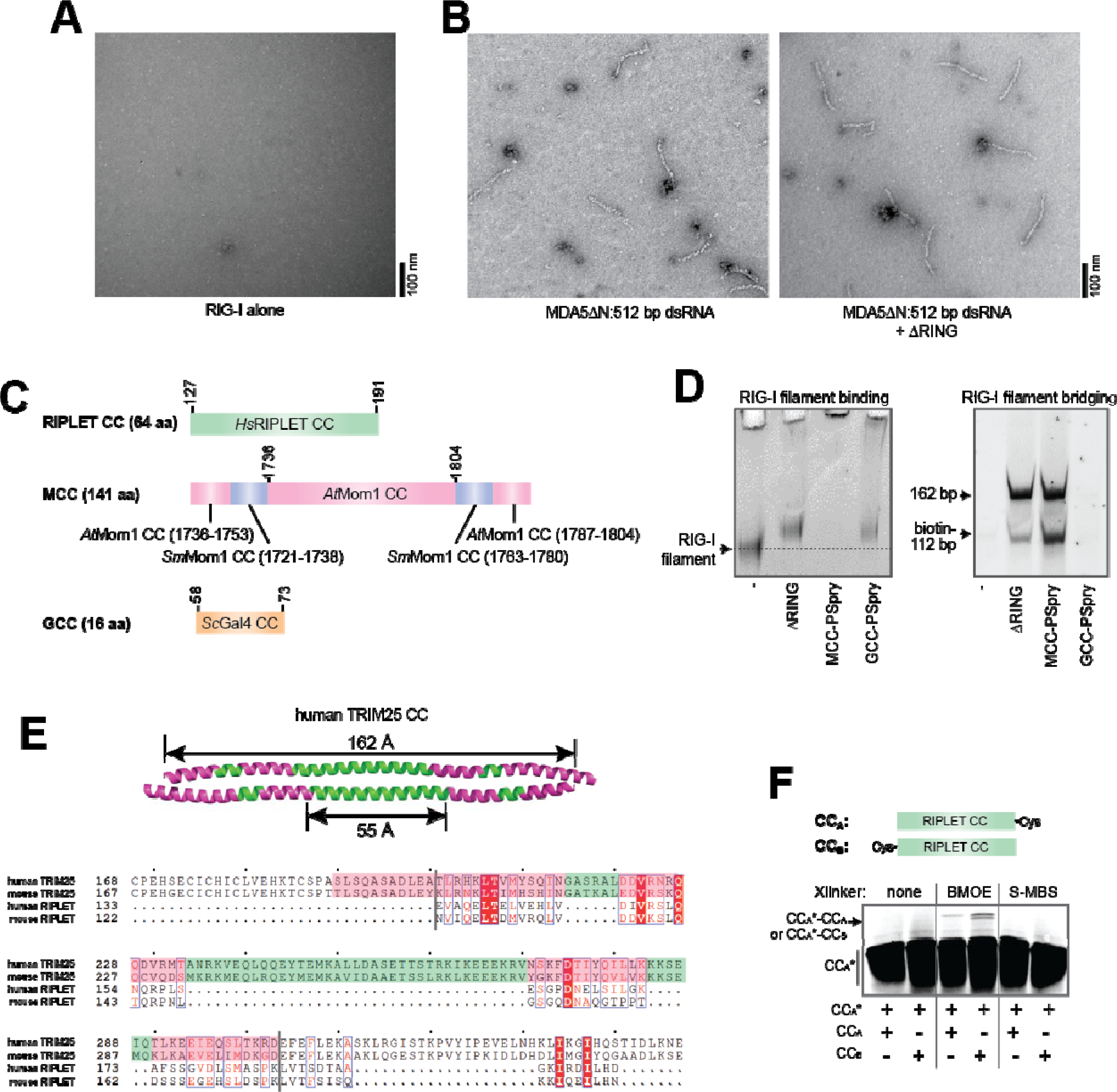
RIPLET cross-bridges RIG-I filaments in an Ub-independent manner. a. Representative EM image of RIG-I protein alone (0.5 μM).
b. Representative EM images of MDA5ΔN filaments (0.4 μM) formed on 512 bp dsRNA (1 ng/μl), in the presence and absence of ΔRING (0.4 μM). Due to the instability of the MDA5 filament on EM grids, ADP•AlF_4_ (1 mM) was added prior to the grid preparation. Previous EM analysis of MDA5 filaments similarly required ADP•AlF_4_ to stabilize the filament on thhee grid (Peisley et al., 2011; Wu et al., 2013).
c. Schematic of coiled coils (CCs) used in this study. *Homo sapiens* RIPLET CC (*Hs*RIPLET CC) indicates residues 127-191. <u>*Saccharomyces cerevisiae*</u> Gal4 CC (*Sc*GCC) indicates residues 58-73. Mom1 CC (MCC) was generated by symmetrically extending *Arabidopsis thaliana* Mom1 CC (*At*Mom1, residues 1736-1804) (Nishimura et al., 2012) with parts of *At*Mom1 CC and *Selaginella moellendorffii* MOM CC (*Sm*Mom1 CC) to generate 141 amino acid-long antiparallel CC.
d. Comparison of CC variants of RIPLET in terms of RIG-I filament binding and cross-bridging activities. Left: Native gel shift assay showing that RIG-IΔN filament (0.5 μM) on 512 bp dsRNA (1 ng/μl) is efficiently bound by ΔRING, MCC-PSpry and GCC-PSpry (0.5 μM for all). Although RIG-IΔN filaments in complex with MCC-PSpry do not enter the gel, the complex formation is evident by the complete depletion of free RIG-IΔN filaments upon addition of MCC-PSpry. Right: Streptavidin pull-down analysis to examine RIG-I filament cross-bridging by ΔRING, MCC-PSpry and GCC-PSpry. The experiment was performed as in Figure 4C. The result shows that ΔRING and MCC-PSpry can cross-bridge RIG-I filaments, while GCC-PSpry cannot. Note that purification of even the biotinylated 112 bp dsRNA was dependent on ΔRING (or MCC-PSpry). This result suggests that biotinylated 112 bp dsRNA is also purified largely through the filament cross-bridging mechanism, rather than through direct binding to the streptavidin beads.
e. Sequence analysis of the RIPLET coiled coil (CC) domain. NCBI Blast analysis of RIPLET identifies TRIM25 as the most closely related protein (sequence identity 32%, and 61% homology) among the human genes (UniProt database). Bottom: Sequence alignment of RIPLET CC and TRIM25 CC using Clustal Omega (EMBL-EBI). Top: The structure of TRIM25 CC (PDB code: 4LTB)(Sanchez et al., 2014). As with other members of the TRIM family characterized to date (Sanchez et al., 2014; Weinert et al., 2015), TRIM25 CC displays the antiparallel dimeric geometry. The TRIM25 CC was colored according to its sequence homology to RIPLET CC (green and magenta for aligned and non-aligned regions, respectively). The symmetric arrangement of the homologous regions suggests that RIPLET CC is also likely to be antiparallel. This notion was further supported by the cross-linking analysis in (F).
f. Cys-Cys crosslinking to distinguish between parallel and antiparallel CC geometries. RIPLET CC with Cys at either the C-terminus (CC_A_) or N-terminus (CC_B_) was purified as a homo-dimer. CC_A_ was N-terminally labeled with fluorescein (CC_A_*) and denatured and refolded with an equimolar amount of unlabeled CC_A_ or CC_B_. The refolded dimer was cross-linked using a Cys-Cys specific crosslinker, BMOE (100 μM), or a Cys-Lys specific crosslinker, S-MBS (100 μM, control). Dimer cross-linking was analyzed by SDS-PAGE using fluorescein fluorescence. The more prominent crosslinking between CC_A_* and CC_B_ than between CC_A_* and CC_A_ suggests that RIPLET CC is likely an antiparallel dimer. All data are representative of three independent experiments.

**Supplementary Fig. 5.**
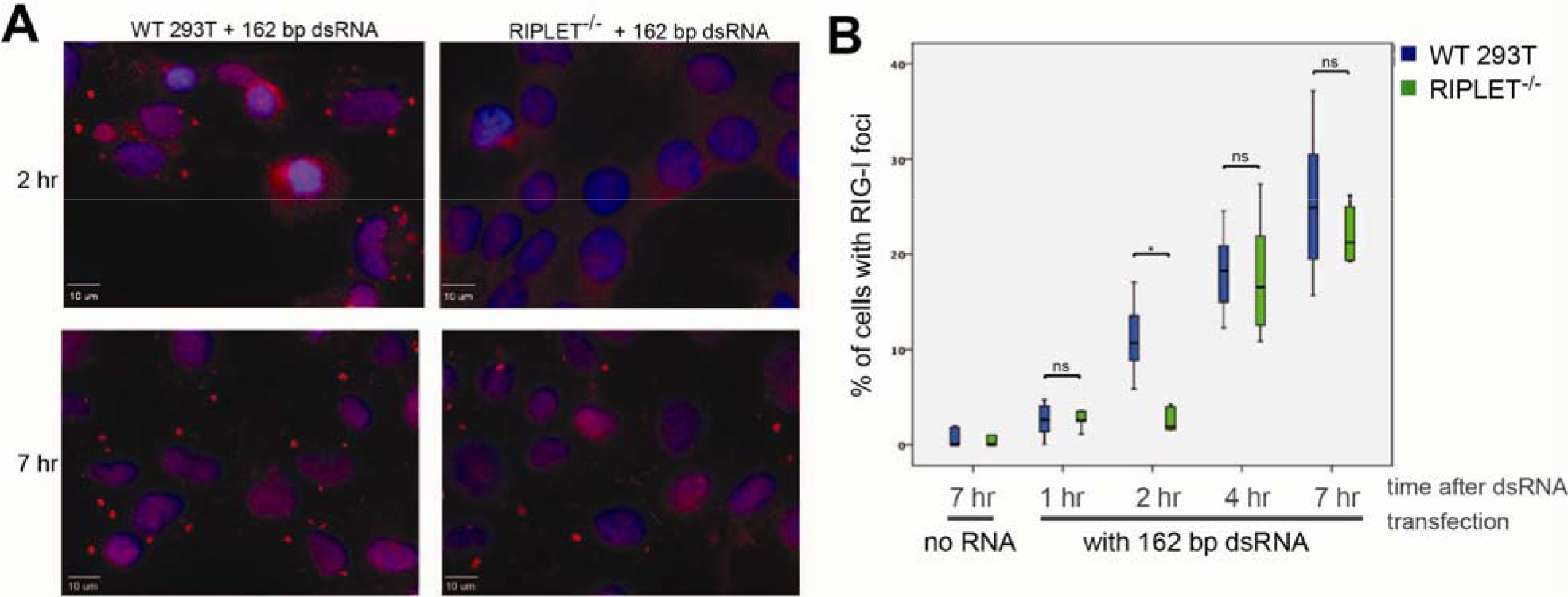
RIG-I localizes at stress granules (SGs) upon dsRNA stimulation, and RIPLET contributes to the kinetics of RIG-I accumulation at SGs, not the steady state level. A. Immunofluorescence analysis of endogenous RIG-I SGs in 293T cells (WT and RIPLET^−/−^) upon stimulation with 162 bp dsRNA (0.5 μg). Cells were fixed 2 hrs or 7 hrs post dsRNA transfection.
B. Quantitation of RIG-I SG assembly kinetics in WT and RIPLET^−/−^ 293T cells. Cells were fixed at indicated time points post RNA transfection and percentage of cells displaying RIG-I SGs were counted using αRIG-I stained images. About 300 cells were examined for each time point per sample. This result suggests that the observed effect of RIPLET on RIG-I SG (Oshiumi et al., 2013) reflects the kinetic delay, not the steady state RIG-I SG level. The delay in RIG-I SG formation can be explained by RIPLET-mediated filament bridging, which likely enables more rapid accumulation of RIG-I in SGs. * *p* < 0.05, ** *p* < 0.01, \*\*\**p* < 0.001, ns, not significant (unpaired *t* test). All data are representative of three independent experiments.

**Supplementary Fig. 6.**
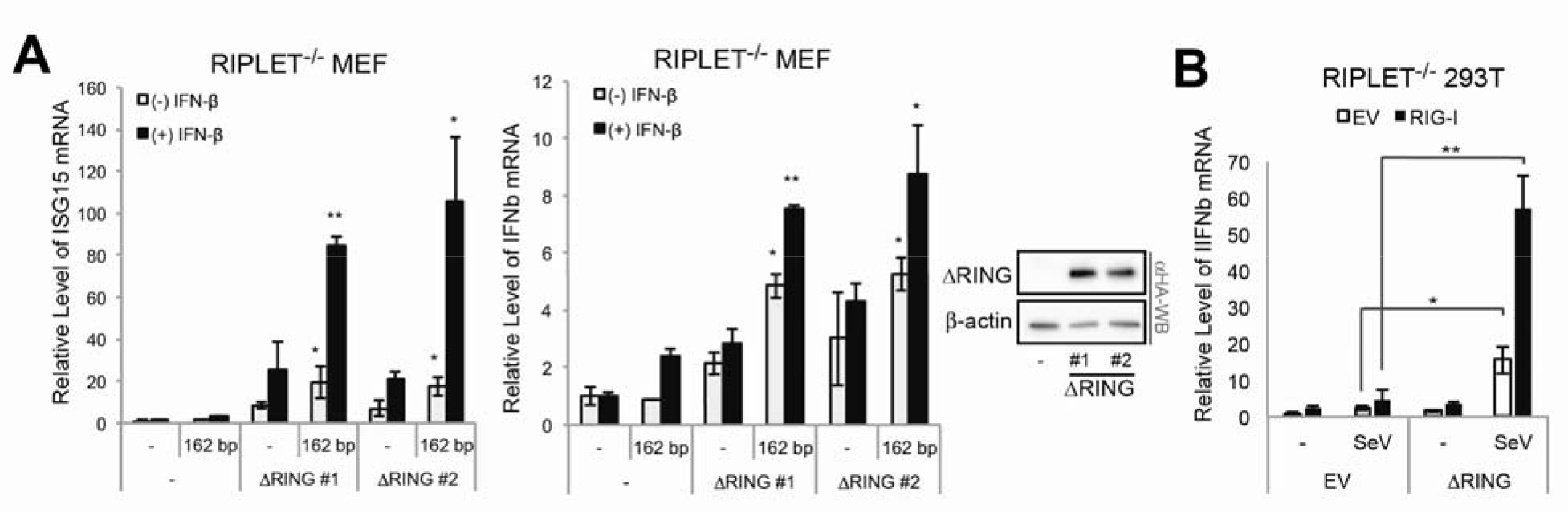
RIPLET-mediated RIG-I aggregation enhances antiviral signaling. a. Relative level of antiviral signaling (mean ± s.d., *n* = 2) in RIPLET^−/−^ MEFs stably expressing ΔRING upon stimulation with 162 bp dsRNA (0.2 μg). Two different clones of RIPLET^−/−^ MEF reconstituted with HA-tagged ΔRING were used (see WB on the right). In a subset of samples, IFNβ (10 ng/ml) was added to the medium (24 hrs prior to RNA transfection) to increase the level of RIG-I. The RIG-I-stimulatory activity of ΔRING was evident even with the basal level of RIG-I and stably expressing ΔRING, but IFNβ further enhanced the effect of ΔRING. This is consistent with the notion that the higher level of RIG-I allows more efficient filament formation on dsRNA and thus, more efficient filament bridging by ΔRING. See also (B).
b. Relative level of antiviral signaling (mean ± s.d., *n* = 3) in RIPLET^−/−^ 293T upon stimulation with 162 bp dsRNA (0.2 μg), with and without complementation with ΔRING. ΔRING was transiently expressed (20 ng) together with RIG-I (10 ng) or by itself in RIPLET^−/−^ 293T. The RIG-I-stimulatory activity of ΔRING was evident even with endogenous RIG-I, but ectopic expression of RIG-I further enhanced the effect of ΔRING. All data are representative of three independent experiments. * *p* < 0.05, ** *p* < 0.01, \*\*\**p* < 0.001, ns, not significant (unpaired *t* test, compared to no ΔRING).

## Methods and Materials

### Key resources table

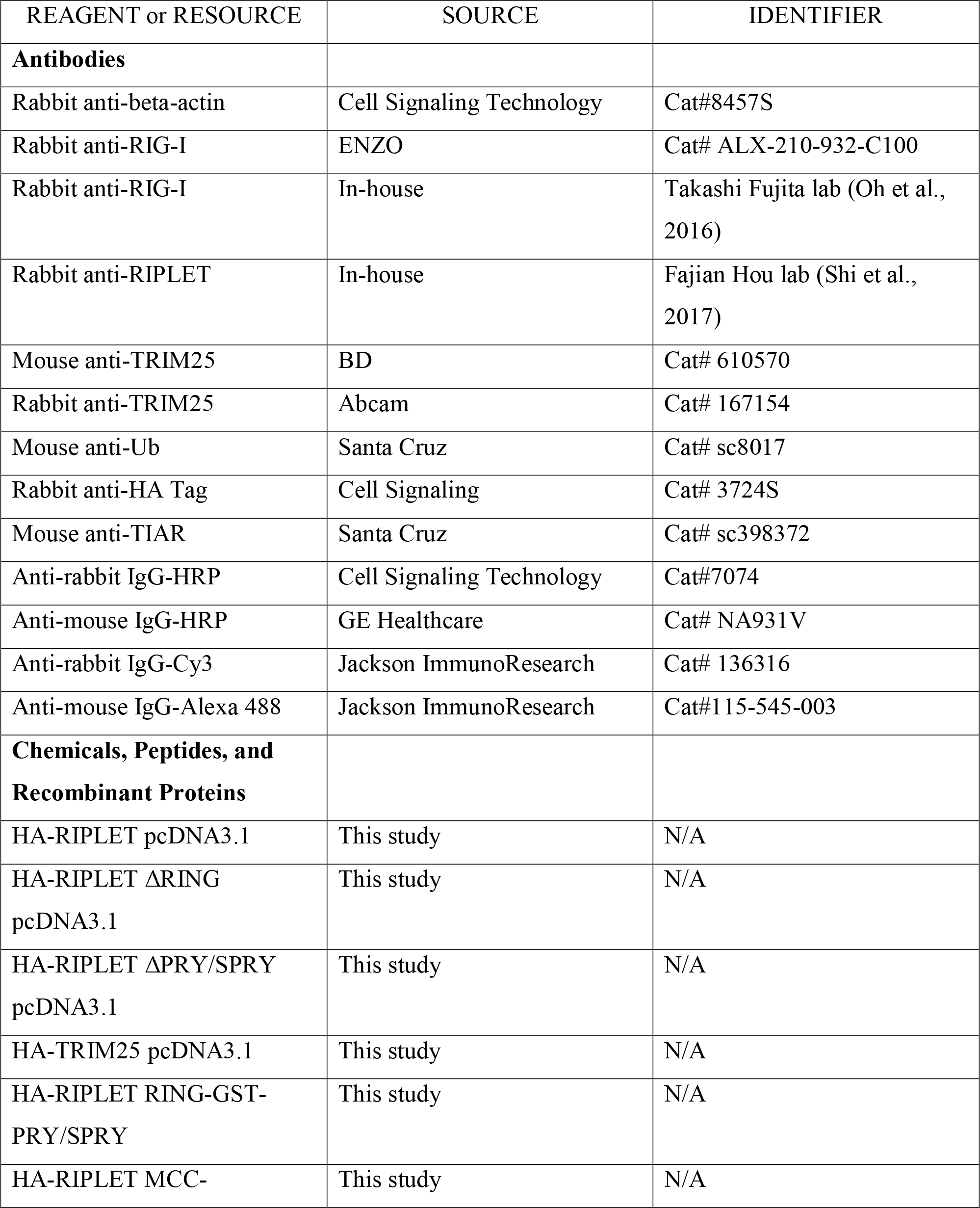

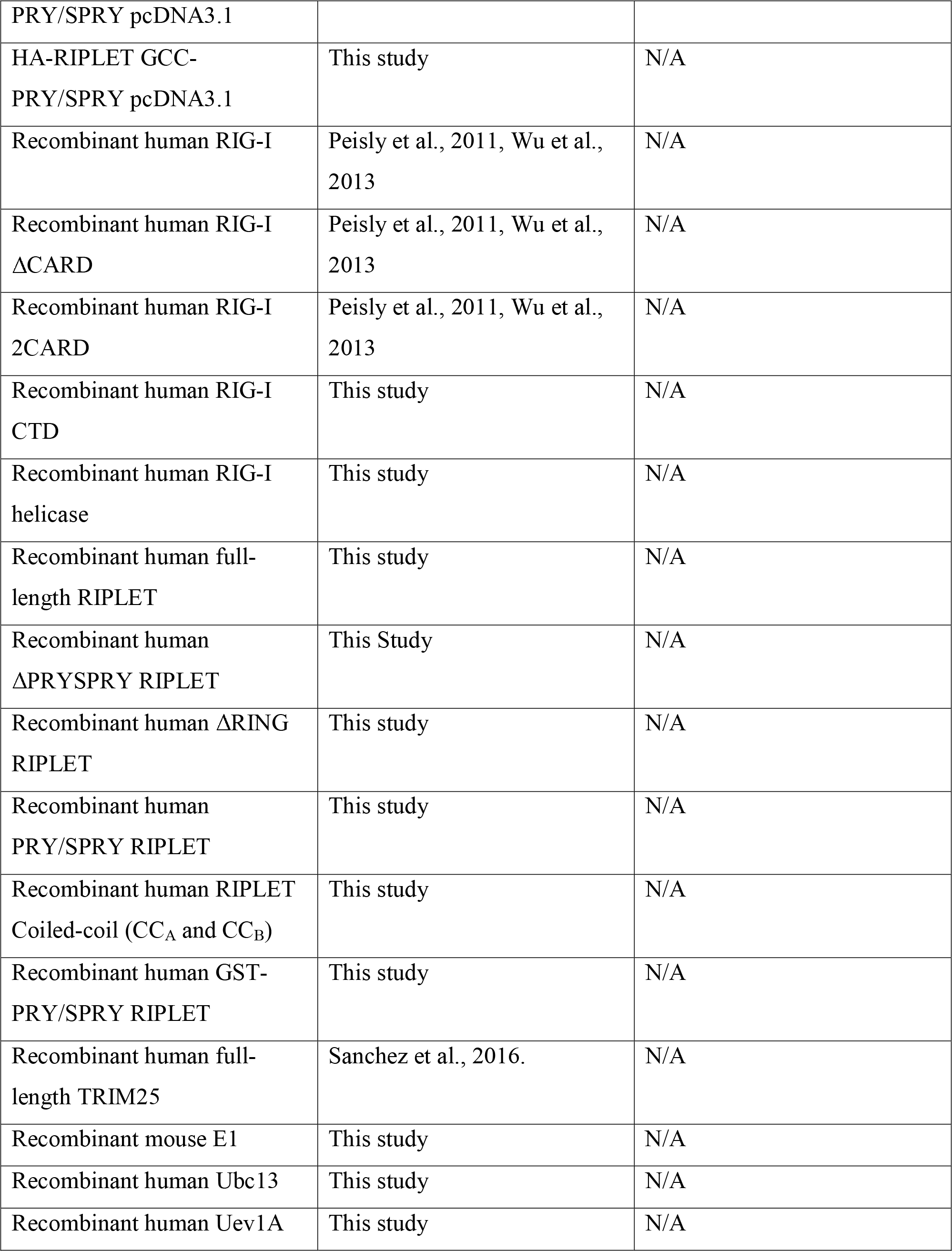

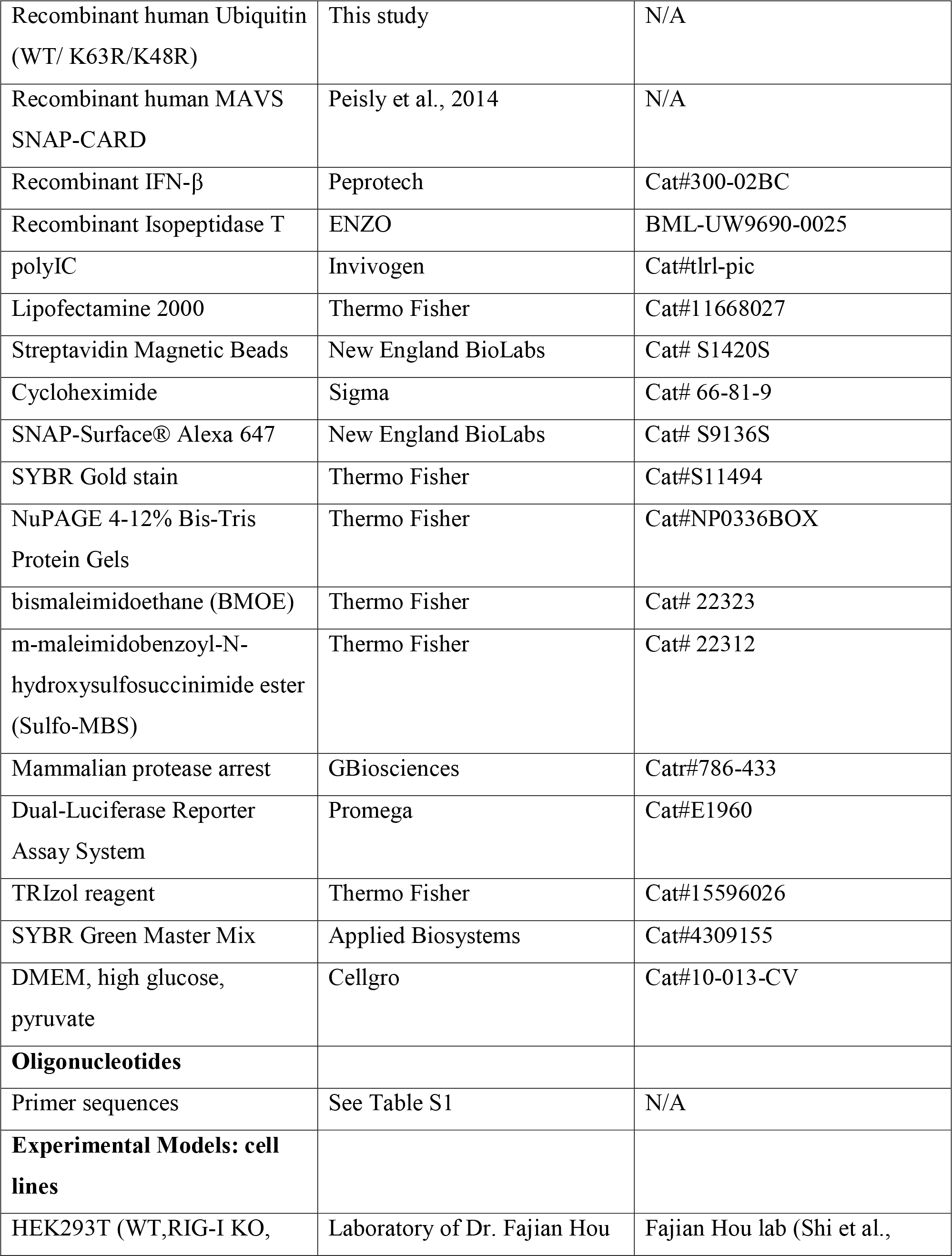

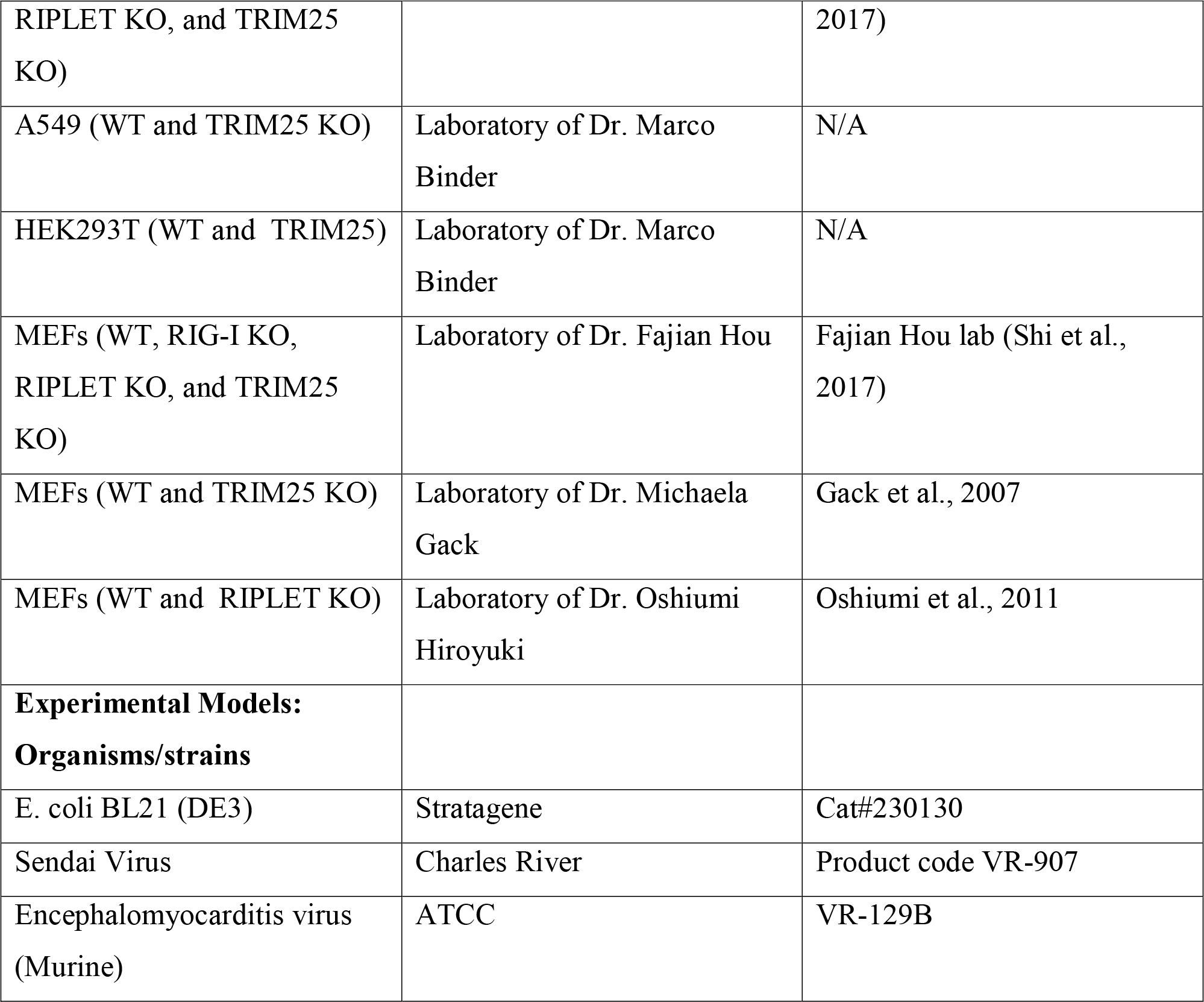

### Contact for reagent and resource sharing

Further information and requests for reagents may be directed to, and will be fulfilled by the corresponding author Sun Hur.

## Experimental Model and Subject Details

### Cell Lines

#### HEK293T cells

Cells were maintained in DMEM (High glucose, L-glutamine, Pyruvate) with 10% fetal bovine serum, 1% penicillin/streptomycin.

#### A549 cells

Cells were maintained in DMEM (High glucose, L-glutamine, Pyruvate) with 10% fetal bovine serum, 1% penicillin/streptomycin.

#### MEF cells

Cells were maintained in DMEM (High glucose, L-glutamine, Pyruvate) with 10% fetal bovine serum, 1% penicillin/streptomycin. MEF cell lines stably expressing RIPLET were additionally maintained with 1mg/mL Neomycin.

### Material Preparation

#### Cell lines

Unless mentioned otherwise WT, RIG-I^−/−^, RIPLET^−/−^ and TRIM25^−/−^ 293T and MEF cell lines generated by the CRISPR/Cas9 technology from the Hou lab (Shi et al., 2017) were used throughout the manuscript. Additional RIPLET^−/−^ and parental WT MEF cell lines in Figure S1 were obtained from the Oshiumi lab (Oshiumi et al., 2010). Also used in this study (Figure S1) are TRIM25^−/−^ and parental WT MEFs from the Gack lab (Gack et al., 2007), and TRIM25^−/−^, RIG-I^−/−^ and parental WT 293T and A549 cell lines from the Binder lab. The Binder cell lines were generated by the CRISPR/Cas9 technology using lentivirus transduction and selection with 1 μg/ml puromycin (see ***Viruses***). Monoclonal A549 TRIM25^−/−^ cells were isolated by limiting dilution. Knockout of TRIM25 was validated by Western blot. RIPLET^−/−^ MEFs stably expressing mouse RIPLET or ΔRING were also obtained by lentivirus transduction and selection with 1 μg/ml neomycin (see ***Viruses***). In all cases, parental, passage-matched cell lines were compared for the effect of knock-out or complementation.

#### Plasmids

Mammalian and bacterial expression plasmids for RIG-I, MDA5 and MAVS and their truncation variants were described previously (Peisley et al., 2013; Wu et al., 2013). Human RIPLET, ΔRING (residues 94-433), ΔPSpry (residues 1-237) and PSpry (residues 249-433) were inserted between BamHI and EcoRI of pcDNA3-HA. GST-PSpry was generated first by inserting PSpry (residues 249-443) into pGEX6p-1 between BamHI and SalI. GST-PSpry in pGEX6p-1 was then amplified and inserted into pcDNA3-HA using the EcoRI site (for both 5’ and 3’ end restrictions). ΔRING was mutated into V127L (GTA → CTC) and T189E/S190F (ACTTCC → GAATTC) to engineer XhoI and EcoRI sites, respectively, and the sites were used to replace RIPLET CC by MCC and GCC (Figure S4D). Note that V127L (GTA → CTC) and T189E/S190F (ACTTCC → GAATTC) alone did not impact the function of RIPLET. 4xFKBP-PSpry was generated by inserting four tandem repeats of FKBP (F36V variant, also known as DmrB) into the KpnI site in pcDNA3 containing PSpry between BamHI/EcoRI. Bacterial expression constructs for RIPLET, ΔRING and GCC-PSpry were generated using BamHI and SalI in pET47b. RIPLET ΔPRYSPRY (residues 1-237) was inserted in pET47b using BamHI and EcoRI sites. MCC-PSpry was cloned using BamHI and SalI in pET50b. GST-PSpry was expressed from pGEX6p-1. Human TRIM25 was cloned between EcoRI and XbaI in pcDNA3-HA vector (pcDNA3 with the HA-tag between KpnI and BamHI). Baculovirus expression construct for human TRIM25 was a generous gift from Owen Pornillos’s lab. The bacterial expression construct for mouse E1 ubiquitin activating enzyme (pET28-mE1) was purchased from Addgene. Human Ubc13 and Uev1A were inserted between the XmaI and XhoI restriction sites in pET47b, and human ubiquitin (WT, K63R, K48R) were inserted between the XmaI and HindIII restriction sites in pET47b. Human ubiquitin (K63only) was purchased from AddGene and subcloned into pcDNA3 with the N-terminal His-tag. The gene encoding residues 124 to 186 of RIPLET, corresponding to the coiled coil (CC) region, was inserted in pMAL-3C vector (pMAL where the cleavage site is inserted between KpnI and BamHI(Alexandrov et al., 2001)) between BamHI and HindIII sites. A single Cys residue was incorporated at either end of the coiled coil, separated by a 4 residue linker (Gly-Ser-Gly-Ser), to generate two distinct forms of the construct namely Cys-GSGS-CC and CC-GSGS-Cys.

#### Viruses

SeV (Cantell strain) was purchased from Charles River. VSV (Indiana stain) was a generous gift from Dr. Sean Whelan (Harvard). EMCV (murine) was purchased from ATCC (VR-129B). Rift Valley fever virus expressing *Renilla reniformis* luciferase, in place of the NS genes (RVFVΔNSs_RLuc) (Kuri et al., 2010), was a kind gift from Dr. Friedemann Weber (University of Gießen, Germany), and was propagated in BHK21 cells. Influenza A/SC35M (H7N7) virus carrying the *Gaussia princeps* luciferase (GLuc) on the NS1 segment (FluA_GLuc) (Reuther et al., 2015) was a kind gift from Dr. Martin Schwemmle (University of Freiburg, Germany), and was propagated in DF-1 cells. For knocking-out TRIM25, lenti-CRISPR plasmid (Addgene #52961) encoding TRIM25 specific gRNA (GGTCGTGCCTGAATGAGACG) was generated using the restriction enzyme, BsmBI. 293T cells were transfected with the following three plasmids at a 3:1:3 ratio: (i) pCMV-DR8.91, coding for HIV gag-pol; (ii) pMD2.G, encoding the VSV-G glycoprotein; and (iii) lenti-CRISPR-TRIM25. pCMV-DR8.91 and pMD.2G were kind gifts from Prof. Didier Trono (Lausanne, Switzerland). 48, 56 and 72 h post transfection cell-free supernatants were harvested and used for transduction of target cells. To generate RIPLET^−/−^ MEFs stable cell lines, HA-tagged mouse RIPLET or ΔRING were cloned into pBABE-Neo (kindly provided by Jim DeCaprio) using the restriction sites BamHI and EcoRI. 293T cells were transfected with the following three plasmids: pBABE-Neo, MLV Gag (gift from Dr. James DeCaprio), and pMD2.G VSV-G in a 10:10:1 ratio. Lentiviruses were harvested 48hrs post transfection. RIPLET^−/−^ MEFs (from Oshiumi’s lab) were infected with the 0.45 μM filtered supernatants for 48 hrs, then selected with 1 μg/ml neomycin. Single clones were selected by limiting dilution and confirmed by western blot.

#### Proteins

Human RIG-I and MDA5 variants were expressed and purified as previously reported(Ahmad et al., 2018; Peisley et al., 2013). Briefly, the proteins were expressed in BL21(DE3) at 20°C for 16-20 hr following induction with 0.5 mM IPTG. Cells were lysed by high pressure homogenization using an Emulsiflex C3 (Avestin), and the protein was purified by a combination of Ni-NTA and heparin affinity chromatography and size exclusion chromatography (SEC) in 20 mM Hepes, pH 7.5, 150 mM NaCl and 2 mM DTT. Human RIPLET and its variants were also expressed and purified in the same manner and purified by a combination of Ni-NTA and SEC. Human TRIM25 was expressed from SF9 insect cells and purified as previously described(Sanchez et al., 2016). Mouse E1, human Ubc13 and Uev1A and ubiquitins were prepared as previously reported(Peisley et al., 2013).

For N-terminal fluorescent labeling of RIG-I and its variants, the protein (~2 mg/ml) was incubated with 0.5 mM peptide (LPETGG) conjugated with fluorescein (Anaspec) and 1.5 μM *S. aureus* sortase A (a gift from Hidde Ploegh, MIT)(Antos et al., 2009) at 4°C for 4 hrs, followed by reverse Ni-NTA affinity purification and/or SEC to remove sortase.

MAVS CARD was expressed as a fusion protein with the SNAP tag in BL21(DE3) at 20°C for 16-20 hr following induction with 0.4 mM IPTG. The SNAP tag allows fluorescent labeling of MAVS CARD. MAVS CARD-SNAP fusion was purified using Ni-NTA affinity chromatography and SEC as previously described(Wu et al., 2016), with the exception of using 0.05% NP-40 instead of CHAPS as a choice of detergent. Purified MAVS CARD-SNAP exists in the form of short filaments and was denatured in 6 M guanidinium hydrochloride for 30 min at 37°C, followed by dialysis against 20 mM Tris, pH 7.5, 500 mM NaCl, 0.5 mM EDTA and 20 mM BME at 4°C for 1 hr. Refolded CARD-SNAP was filtered using a 0.1μ filter and subsequently labeled with Alexa647-benzylguanine (NEB) on ice for 15 min according to the manufacturer’s instruction, and was immediately used for filament formation assays.

Both RIPLET CC with Cys at the C-terminus (CC-GSGS-Cys) and N-terminus (Cys-GSGS-CC) were expressed and purified along with a cleavable 6xHis and MBP tag from *E. coli* BL21 (DE3) using Ni-NTA affinity chromatography followed by size exclusion chromatography in (50 mM Tris pH 7.5, 300 mM NaCl, 0.5 mM EDTA). The 6xHis and MBP tags were removed from both proteins using HRV 3C protease. CC-GSGS-Cys was subsequently fluorescently labeled by attaching FAM-conjugated peptide LPETGG (Anaspec) at the N-terminus using the protein ligase Sortase A. Purified CC-GSGS-Cys was incubated with *S. aureus* Sortase A in a molar ratio of 4:1 along with 0.5 mM FAM-LPETGG peptide for 6-8 hrs at RT in the dark followed by reverse Ni-NTA affinity chromatography to remove sortase.

#### RNAs

Double-stranded RNAs (dsRNAs) were prepared by *in vitro* T7 transcription as previously described (Peisley et al., 2011). Unless mentioned otherwise, all dsRNAs contain 5’ppp and blunt end. The sequences of 28, 42, 62, 112, 162 and 512 bp dsRNAs were taken from the first 16, 30, 50, 100, 150 and 500 bp of the MDA5 gene flanked by 5’-gggaga and 5’-tctccc. The sequences of 15 bp and 21 bp dsRNAs are 5’-agggcccgggaugcu and 5’-gagucacgacguuguaaaaaa, respectively. The two complementary strands were co-transcribed, and the duplex was separated from unannealed ssRNAs by 15% acrylamide (15 and 21 bp), 8.5% acrylamide (28-162 bp) or 6% acrylamide (512 bp) gel electrophoresis in TBE buffer. RNA was gel-extracted using the Elutrap electroelution kit (Whatman), ethanol precipitated, and stored in 20 mM Hepes, pH 7.0. Qualities of RNAs were analyzed by 1X TBE polyacrylamide gel.

For 3’-biotin labeling of RNA, the 3′ end of RNA was oxidized with 0.1 M sodium periodate in 0.25 M NaOAc, pH 5.5, at room temperature for 2 h. The reaction subsequently was treated with 0.2 M KCl, the buffer was replaced with 0.25 M NaOAc, pH 5.5, using Zeba desalting column (Pierce), and incubation EZ-Link Hydrazide-PEG4-Biotin (Pierce) continued at room temperature for 4–6 h. The labeled RNA was desalted twice to remove unincorporated biotin and was stored in 20 mM Hepes, pH 7.0, at −20 °C.

#### Antibodies

For immuno-blotting, antibodies and their manufacturers were: Rabbit α-RIG-I (ENZO), rabbit α-HA (Cell Signaling), rabbit α-β-actin (Cells Signaling) mouse α-TRIM25 (BD), mouse α-Ub (Santa Cruz), α-RIPLET (in-house). For immunofluorescence, antibodies included: α-RIG-I (in-house)(Oh et al., 2016) and α-TIAR (Santa Cruz).

### Luciferase reporter assay

For the *IFNβ* promoter reporter assay, HEK293T cells were maintained in 48-well plates in Dulbecco’s modified Eagle medium (Cellgro) supplemented with 10% fetal calf serum and 1% penicillin/streptomycin. At ~80% confluence, cells were transfected with plasmids encoding the *IFNβ* promoter driven firefly luciferase reporter plasmid (100 ng) and a constitutively expressed Renilla luciferase reporter plasmid (pRL-CMV, 10 ng) by using lipofectamine 2000 (Life) according to the manufacturer’s protocol. Unless mentioned otherwise, signaling activity of endogenous RIG-I was measured. In Figure 1D, expression plasmid for RIG-I (10 ng), MDA5 (5 ng), MAVS (25 ng) or GST-2CARD (50 ng), was transfected together with the reporter plasmids. In the experiments requiring RNA stimulation, the medium was changed 6hr after the first transfection and the cells were additionally transfected with RNA (0.2 μg, Invivogen). Cells were lysed ~20 hrs post-stimulation and *IFNβ* promoter activity was measured using the Dual Luciferase Reporter assay (Promega) and a Synergy2 plate reader (BioTek). Firefly luciferase activity was normalized against Renilla luciferase activity.

For RVFV and FluA luciferase reporter assay in Figure S1D-S1E, cells were infected with RVFVΔNSs_RLuc (MOI=0.05) or FluA_GLuc (MOI=0.001) for one hour in Opti-MEM, before the medium was exchanged with fresh DMEM (2% FCS). Cells were lysed at indicated time points using Luc-lysis buffer (1% Triton X-100, 25 mM glycyl-glycin, pH 7.8, 15 mM MgSO_4_, 4 mM EGTA, 10% glycerol). 100 μl of cell lysate (for RVFVΔNSs_RLuc) or 100 μl of medium supernatant (for FluA_GLuc) was mixed with 400 μl of luciferase assay buffer (15 mM K_3_PO_4_ [pH7.8], 25 mM glycyl-glycin [pH 7.8], 15 mM MgSO_4_, 4 mM EGTA) supplemented with 1.43 μM coelenterazine (PJK, Germany). The luciferase buffer was injected into each well before measurement using a Mitras^2^ multimode plate-reader LB942 (Berthold, Germany) and the luciferase activity was measured using a 480 nm filter (480m20BREThs, Berthold). Upon luciferase measurement of each well, reaction was quenched with 100 μl of 10% SDS to prevent bleed-through into the neighboring wells.

### RT-qPCR

Unless mentioned otherwise, endogenous RIG-I signaling activity was measured by measuring the level of mRNAs for IFNβ or indicated IFN-stimulated genes. HEK293T cells were stimulated with dsRNA in a similar manner in the luciferase assay. For SeV infection, cells were treated with SeV (100 HA units/ml, Charles River) with 1 hr of absorption. For EMCV infection, MEFs were treated with the virus at a MOI = 0.1 with 1 hour absorption. Cells were lysed ~20 hr post-infection. Total RNAs were extracted using TRIzol reagent (Thermo Fisher) and cDNA was synthesized using High Capacity cDNA reverse transcription kit (Applied Biosystems) according to the manufacture’s instruction. Real-time PCR was performed using a set of gene specific primers (see the Table S1), a SYBR Green Master Mix (Applied Biosystems), and the StepOne™ Real-Time PCR Systems (Applied Biosystems). For RVFV and FluA infection, cells were infected with RVFVΔNSs_RLuc (MOI=0.05) or FluA_GLuc (MOI=0.001) as described above. Cells were harvested ~20 hr or ~40 hr post-infection, and RNA was isolated using the NucleoSpin RNA Kit (Macherey-Nagel) according to manufacturer’s protocol. cDNA was generated as described above and real-time PCR was performed using SYBR Green (Biorad) and gene-specific primers (see the Table S1) using the CFX96 real-time system (Biorad).

### Ubiquitination reaction

RIG-I ubiquitination was performed in the following manner. Purified RIG-I (1.0 μM) was first incubated with dsRNA (2 ng/μl) in buffer A (20 mM Hepes, pH 7.5, 150 mM NaCl, 1.5 mM MgCl_2_, 2 mM ATP and 5 mM DTT) at RT for 15 min. The RIG-I:RNA complex (to the final RIG-I concentration of 0.5 μ M) was then further incubates with 20 μM ubiquitin, 1 μM mE1, 5 μM Ubc13, 2.5 μM Uev1A and 0.25 μM E3 (RIPLET or TRIM25) in buffer B (50 mM Tris pH 7.5, 10 mM MgCl_2_, 5 mM ATP and 0.6 mM DTT) at 37°C for 30 min. For isoT treatment, isoT (10 ng/μl final) was added to the ubiquitination reaction, and further incubated at 37°C for 30 min. Reactions were quenched with SDS loading buffer, boiled at 96 C for 5 min, and analyzed on SDS-PAGE by anti-RIG-I or anti-Ub western blot. Note that previous Ub-conjugation of RIG-I CARD by TRIM25 (Peisley et al., 2014) was performed with significantly higher protein concentrations, which explains the discrepancy with the current result in Figure 2.

Unanchored K63-Ub_n_ chains were generated from a reaction containing 0.4 mM ubiquitin, 4 μM mE1, 20 μM Ubc13 and 20 μM Uev1A in buffer B. The K63-Ub_n_ synthesis reaction was performed overnight at 37°C. Synthesized K63-Ub_n_ chains were purified as described previously (Dong et al., 2011). Briefly, ubiquitin chains were diluted 5-fold into 50 mM ammonium acetate, pH 4.5, 0.1 M NaCl and separated over a 45 ml 0.1-0.6 M NaCl gradient in 50 mM ammonium acetate, pH 4.5 using a Hi-Trap SP FF column (GE Healthcare). High molecular weight fractions were applied to an S200 10/300 column equilibrated in 20 mM Hepes pH 7.5, 150 mM NaCl.

### MAVS polymerization assay

The MAVS filament formation assay was performed as previously reported (Wu et al., 2013). Refolded, Alexa647 labeled monomer of CARD fused to SNAP (CARD-SNAP) was described in the material preparation section. In the absence of external stimuli or seed filaments, refolded MAVS CARD remains stably as a monomer over ~10-20 hr, after which it spontaneously forms prion-like filaments over the course of days. Thus, all assays involving MAVS filament formation were performed within 4 hr after refolding. To monitor stimulation of MAVS filament formation by RIG-I conjugated with ubiquitin chains, monomeric MAVS CARD-SNAP (10 μM) was incubated with conjugation reaction (to final RIG-I concentration of 1 μM) at RT for 1 hr prior to analysis by Bis-Tris native PAGE (Life) or by EM. Before running on the gel, all samples were subjected to one round of freeze-thaw cycle by incubating on dry ice for 5 min followed by incubation at RT for 5 min. To examine the impact of ΔRING on MAVS stimulatory activity of RIG-I, RIG-I (1 μM) was pre-incubated with dsRNA (6 ng/μl) and 2 mM ATP in the presence or absence of K63-Ub_n_ (equivalent to 0-7 μM monomeric Ub) in buffer A for 15 min at RT. This was followed by the addition of ΔRING (0, 5, 12.5 μM) and a further incubation at RT for 15 min. The mixture was then finally incubated with monomeric CARD-SNAP (10 μM) at RT for 1 hr prior to analysis by Bis-Tris native PAGE (Life) or by EM. Fluorescent gel images were recorded using an FLA9000 scanner (Fuji).

### Electron Microscopy

RIG-I or RIG-IΔN (0.5 μM) was incubated with RNA (1 ng/μl) in buffer A at RT for 15 min followed by addition of RIPLET, ΔRING or its CC-swap variants (0.5 μM) and further incubated at RT for 15 min prior to EM grid preparation. Prepared samples were adsorbed to carbon-coated grids (Ted Pella) and stained with 0.75% uranyl formate as described(Ohi et al., 2004). Images were collected using a Tecnai G^2^ Spirit BioTWIN transmission electron microscope at 30,000x magnification for RIG-I filaments and 4,800x or 6,800x for MAVS filaments. Grids for the MAVS filaments were prepared and imaged as with the RIG-I filaments.

### Native gel shift assays

For RNA binding assays, dsRNA (1 ng/μl) was incubated with protein (at indicated concentration) in buffer A at RT for 10 min, and the complex was analyzed on Bis-Tris native PAGE (Life Technologies) after staining with SybrGold stain (Life Technologies). For RIG-I binding assays, fluorescein-labeled RIG-I was first incubated with dsRNA (1 ng/μl) and subsequently with RIPLET (at indicated concentration) prior to analysis by Bis-Tris native PAGE (Life Technologies). SybrGold or fluorescein fluorescence was recorded using the scanner FLA9000 (Fuji) and analyzed with Multigauge (GE Healthcare).

### Streptavidin pull-down assay

RIG-IΔN filaments (0.5 μM) were formed on non-biotinylated 162 bp dsRNA (1 ng/μl) and 3’-biotinylated 112 bp dsRNA (1 ng/μl), and were incubated with an increasing concentrations of ΔRING or GST-PSpry (0.25, 0.5, and 1μM) at RT for 15 min in buffer A. The reaction (250 μl) was further incubated with streptavidin-agarose (15 μl, NEB), washed three times with 0.5 ml buffer A, and eluted with 1 unit/ml protease K (NEB). Biotinylated 112 bp dsRNA was purified using streptavidin beads and co-purification of 162 bp dsRNA was analyzed on the 6% TBE gel. Gel image was taken using SybrGold stain fluorescence.

For cellular streptavidin pull-down experiments, WT and RIPLET^−/−^ 239T cells were treated with 100 μg/mL cyclohexamine for 1 hr, and then transfected with non-biotinylated 162 bp dsRNA (1 μg) and 3’ biotinylated 112 bp dsRNA (1 μg). Cells were harvested 4 hrs post-transfection in the hypotonic lysis buffer (20 mM Tris, pH 7.5, 10 mM KCl, 1.5 mM MgCl_2_, 250 mM sucrose) and lysed using a 25G syringe (5X). Supernatants were cleared first at 500g for 4 minutes, then at 10000g for 4 minutes. The cleared lysate (400 μl) was incubated with 50 μl streptavidin-magnetic beads (NEB) overnight. The beads were washed with 500 μl of hypotonic lysis buffer (2X), 500 μl of wash buffer (0.5M NaCl, 20mM Tris-HCl, pH 7.5, 1mM EDTA), and 200 μl of ice-cold low salt buffer (0.15M NaCl, 20mM Tris-HCl, pH 7.5, 1mM EDTA). RNA was eluted in 30 μl of elution buffer (10mM Tris-HCl (pH 7.5), 1mM EDTA) at 65°C for 5 min, then purified using TRIzol reagent. cDNA was synthesized using High Capacity cDNA reverse transcription kit with 162bp dsRNA-specific primers, and qPCR was performed as described above.

### Immunofluorescence imaging

293T cells were infected with SeV or stimulated with 162 bp dsRNA with 5’ppp. At indicated time points post RNA stimulation, cells were fixed with 2% paraformaldehyde at RT for 10 minutes, and permeabilized with 0.5% Triton-X at RT for 10 minutes. Cells were blocked for 30 min at RT with 1% BSA in PBST, and stained using RIG-I (1:100, in-house antibody) and TIAR (1:100, SC-398372) overnight. Images were obtained using a Zeiss Axio Imager M1 microscope at 100X magnification.

### Multi-Angle Light Scattering (MALS)

The molecular masses of RIPLET, ΔRING and PSpry were determined by MALS using a Superose 200 10/300 column (GE) attached to a miniDAWN TRI-STAR detector (Wyatt Technology) in 20 mM Hepes, pH 7.5, 150 mM NaCl, 2 mM DTT and the data was analyzed using Astra V (Wyatt Technology).

### Coiled coil cross-linking assay

The fluorescently labeled CC-GSGS-Cys (CC_A_* in Figure S4F) was mixed with equal molar amounts of unlabeled CC-GSGS-Cys (CC_A_) or Cys-GSGC-CC (CC_B_) in denaturation buffer (50 mM Tris 7.5, 300 mM NaCl, 0.5 mM EDTA, 20 mM BME, 6 M Urea) and incubated at 37°C for 30 min with constant shaking. The denatured protein mixtures were refolded by stepwise dilution into refolding buffer (denaturation buffer without Urea and BME). At each step, the mixture was diluted two-fold into refolding buffer and incubated at 4°C for 15 min with gentle rocking and the step repeated 4 times. Finally the refolded proteins were desalted into refolding buffer using 7K MWCO Zeba spin desalting columns (Thermo Fisher Scientific).

The refolded protein dimers were cross-linked by incubating with 100 μM crosslinker bismaleimidoethane (BMOE) or m-maleimidobenzoyl-N-hydroxysulfosuccinimide ester (Sulfo-MBS) (Thermo Fisher Scientific) at RT for 10 min. The cross-linked dimers were run on a 20% SDS-PAGE and scanned for FAM fluorescence using FLA9000 scanner (Fuji).

### Quantification and Statistical analysis

Average values and standard deviations were calculated using Microsoft excel and SPSS (IBM). The values for *n* represent biological replicates for cellular experiments or individual samples for biochemical assays. For immunofluorescence experiments, *n* represent the number of cells counted. About 300 cells were examined for each time point per sample. For each figure, the number of replicates is indicated in the figure legends. Unless otherwise mentioned, cell culture assays were performed in 3 independent experiments. *p* values were calculated using the two-tailed unpaired Student’s t test. We consider values * *p* < 0.05, ** *p* < 0.01, \*\*\**p* < 0.001 significant.

